# Mechanism of Proton Release during Water Oxidation in Photosystem II

**DOI:** 10.1101/2024.07.03.602004

**Authors:** Friederike Allgöwer, Maximilian C. Pöverlein, A. William Rutherford, Ville R. I. Kaila

## Abstract

Photosystem II (PSII) catalyzes the light-driven water oxidation that releases dioxygen into our atmosphere and provides the electrons needed for the synthesis of biomass. The catalysis occurs in the oxygen-evolving oxo-manganese-calcium (Mn_4_O_5_Ca) cluster that drives the stepwise oxidation and deprotonation of substrate water molecules leading to the O_2_ formation. However, despite recent advances, the mechanism of these reactions remains unclear and much debated. Here we show that the light-driven Tyr161_D1_ oxidation adjacent to the Mn_4_O_5_Ca cluster, significantly decreases the barrier for proton transfer from the putative substrate water molecule (W3/W_x_) to Glu310_D2_, which is accessible to the luminal bulk. By combining hybrid quantum/classical (QM/MM) free energy calculations with atomistic molecular dynamics (MD) simulations, we probe the energetics of the proton transfer along the Cl1 pathway. We demonstrate that the proton transfer occurs via water molecules and a cluster of conserved carboxylates, driven by redox-triggered electric fields directed along the pathway. Glu65_D1_ establishes a local molecular gate that controls the proton transfer to the luminal bulk, whilst Glu312_D2_ acts as a local proton storage site. The identified gating region could be important in preventing back-flow of protons to the Mn_4_O_5_Ca cluster. The structural changes, derived here based on the dark-state PSII structure, strongly support recent time-resolved XFEL data of the S_3_→S_4_ transition (Nature **617**, 2023), and reveal the mechanistic basis underlying deprotonation of the substrate water molecules. Our combined findings provide insight into the water oxidation mechanism of PSII and show how the interplay between redox-triggered electric fields, ion-pairs, and hydration effects control proton transport reactions.

**Significance Statement:** Photosystem II is nature’s water splitting enzyme that produces the oxygen in the atmosphere and drives the synthesis of biomass. The water splitting reaction releases protons to the luminal bulk contributing to the protonmotive force that drives the synthesis of ATP. Key mechanistic principles of the light-driven water splitting reaction remain debated, amongst them the catalytically important deprotonation steps. Here we show how the oxygen-evolving oxo-manganese-calcium cluster transports protons via conserved carboxylates and water molecules in proton arrays that lead to the luminal bulk. We identify a local proton storage site and molecular gates that prevent wasteful back reactions by undergoing conformational changes, and we show how electric field effects control the protonation dynamics in Photosystem II.

## Introduction

The light reactions of photosynthesis transduce the energy of photons into redox energy and an electrochemical proton gradient across the thylakoid membrane, providing the reducing power for the synthesis of biomass and the driving force for the synthesis of ATP (1, 2). Photosystem II (PSII) functions as the light-driven water/plastoquinone oxidoreductase, which drives electron transfer from water to plastoquinone, while the protons released and taken up at the respective donor and acceptor sides establish a proton motive force (*pmf*) across the photosynthetic membrane. PSII comprises 37 light-capturing chlorin pigments, including a central P_D1_/P_D2_ chlorophyll pair, which has one of the highest oxidation potentials known in biology (1.2-1.4 V) (3, 4). Absorption of light energy triggers a charge separation forming a Chl_D1+_Pheo_D1-_ pair followed by electron transfer from P_D1_ to Chl_D1+_ and from Pheo_D1-_ to Q_A_ and further to Q_B_ (Figure 1A) (5). The cationic P_D1•+_ is reduced by a redox-active tyrosine residue (Tyr161_D1_, Y_z_), forming a tyrosyl radical that oxidizes the oxygen-evolving oxo-manganese calcium (Mn_4_O_5_Ca) (6, 7). The transient formation of the tyrosyl radical (Y_z_^•^) involves a proton-coupled electron transfer (PCET) process that moves the phenolic proton to its hydrogen-bonded base, His190_D1_ (Figure 1C) (8). Two light flashes sequentially reduce Q_B_ to Q_B_H_2_, with a proton uptake from the bulk taking place in each step. Q_B_H_2_ is released from PSII into the thylakoid membrane, carrying the redox energy to the photosynthetic *b*_6_*f* complex (9). Each oxidation of the Mn_4_O_5_Ca releases one proton per electron on average, although at functional pH values, the S_1_ → S_2_ step is not linked to deprotonation, while the combined S_3_→S_4_→S_0_ steps release two sequential protons (10). Four excitations take the Mn_4_O_5_Ca through the S_0_ → S_4_ intermediates of the enzyme cycle (11), which result in the subsequent oxidation of two substrate water molecules into molecular dioxygen (Figure 1C). Despite recent structural advances (12-17), particularly from time-resolved structures obtained using the x-ray free electron laser (XFEL) (18-22), the exact water oxidation mechanism still remains unsolved.

**Figure 1.**
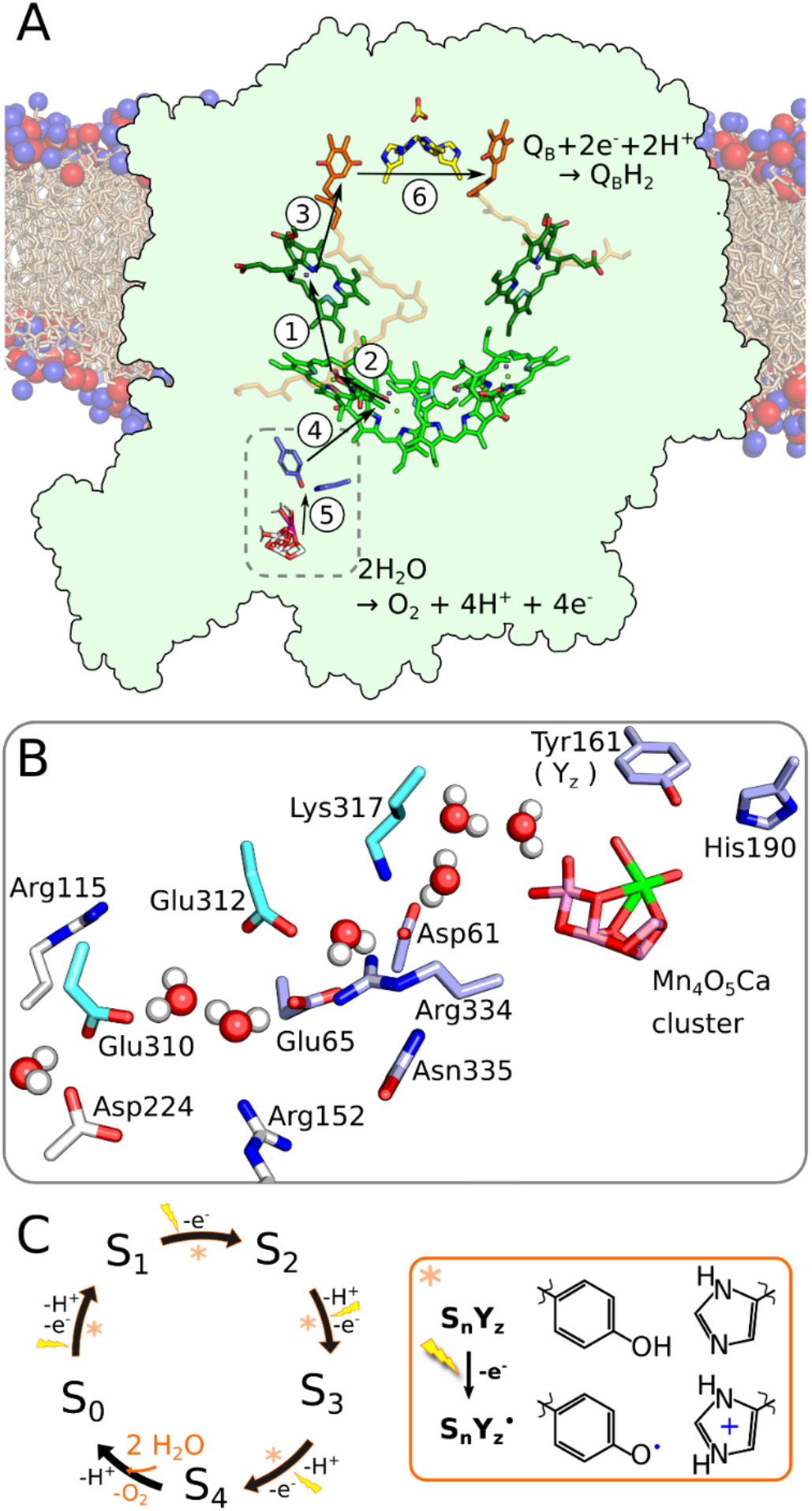
Structure and function of PSII. **A)** Photosystem II from *T. vulcanus* (PDB ID: 3WU2 (12)). The order of the electron transfer steps is indicated by the numbered arrows. **B)** Proton transfer between the Mn_4_O_5_Ca cluster and the luminal bulk, with important residues highlighted (D1 in light blue, D2 in cyan, and PsbO in white). **C)** The stepwise oxidation of the Mn_4_O_5_Ca cluster along the Kok cycle. *Inset:* Each transition leads to a radical formation by oxidation of tyrosine Z (Y_z_) and proton transfer to His190, forming Tyr_Z_^•^/HisH^+^.

Allgöwer *et al*. (23) recently suggested that the Ca^2+^-bound water molecule, W3, deprotonates stepwise and moves to Mn1 upon the S_2_→S_3_ transition (cf. also Refs. (24),(25)). Further deprotonation of this putative substrate water leads to formation of an oxyl radical that can form an O-O bond with the bridging O5 ligand with a low kinetic barrier (23, 24). Although the origin of the proposed substrate water molecule was different, a similar radical-based O-O formation mechanism was also suggested by Siegbahn (26, 27), while acid-base mechanisms have also been proposed (28, 29) [but cf. ref. (30)]. It was also suggested (23) that the formation of the Y_z_^•^/HisH^+^ pair creates an electric field that favors the proton transfer to Asp61_D1_, with re-arrangement of nearby ion-pairs. In this regard, Bhowmick *et al*. (21) recently observed density for a water or hydroxyl ligand (W_x_) appearing next to the Mn1 site in S_2_→S_3_, indirectly supporting the proposed water-oxidation mechanism (23, 24, 26). Moreover, Greife *et al*. (31) proposed similar redox-triggered re-arrangement of ion-pairs, particularly at Asp61_D1_/Lys317_D2_ [but cf. also (32)], which were suggested to modulate the proton transfer energetics (23).

Three water-filled channels near the Mn_4_O_5_Ca cluster have been resolved based on structural experiments (16, 20, 33, 34) and molecular simulations (23, 35-41). The O1 channel was suggested to deliver substrate water molecules to the Mn_4_O_5_Ca (16, 20, 23), while the O4 and Cl1 channels were suggested to function for the proton release to the bulk (20, 23, 42-44). Conformational changes in the Asp61_D1_/Lys317_D2_ ion-pair linked to proton transfer were recently found in the Cl1 channel (20, 23, 31), which could support that this channel is used for deprotonation of the substrate water molecules. Fourier-transformed infrared spectroscopy (FTIR) experiments in combination with site-directed mutagenesis (45-49) further support the functional relevance of this pathway.

To probe the mechanisms underlying the redox-triggered deprotonation of the substrate water molecules, we combine here microsecond atomistic molecular dynamics (MD) simulations with quantum / classical *ab initio* molecular dynamics (QM/MM-MD) and free energy simulations. Our multi-scale simulations show that the Cl1 channel supports a water-mediated proton release via a chain of carboxylates (Asp61_D1_, Glu312_D2_, Glu65_D1_, Glu310_D2_/Asp224_PsbO_; Figure 1B), with oxidation of the redox-active Y_z_ (coupled to proton transfer to the neighboring His190_D1_) lowering the kinetic barrier for the proton transfer reaction by electric field effects. Our study identifies key gating regions and suggests a general mechanism in which electrostatic fields may control proton transfer reactions in biological energy conversion.

## Results

### The redox-triggered proton transfer from the Mn_4_O_5_Ca cluster to the luminal bulk

To explore the energetics and dynamics of proton release from the Mn_4_O_5_Ca cluster, we performed atomistic molecular dynamics (MD) simulations of PSII embedded in a lipid membrane surrounded by water molecules and ions (see *SI Appendix*, Fig. S1A). These microsecond simulations allowed us to resolve redox-triggered conformational and hydration changes during the proton transfer reactions. Based on our classical MD simulations, we performed density functional theory (DFT)-based QM/MM free energy simulations (QM/MM-US; see *Methods*) and *ab initio*-molecular dynamic simulations (QM/MM-MD; *SI Appendix*, Fig. S1B) that allowed us to further probe the dynamics and energetics of the bond-formation and bond-breaking reactions linked to the proton transfer.

During the *ca*. 10 μs classical MD simulations of PSII, Asp61_D1_, which acts as a proton acceptor for W3/W_x_ in the S_2_ → S_3_ and the S_3_ → S_4_ transition (23), forms a hydrogen-bonded water chain that connects Glu312_D2_, Glu65_D1_, Glu310_D2_, and Asp224_PsbO_ to the luminal bulk (Figure 2) and could support Grotthuss-type proton transfer reactions from the Mn_4_O_5_Ca cluster (see below). We note that the highly conserved Glu65_D1_, Arg334_D1_, and Glu312_D2_ residues form a local dry region in this otherwise well-hydrated pathway (Figure 2A), suggesting that these residues could have a potential gating function controlling the proton release (see below). To probe the proton release mechanism, we next simulated the conformational dynamics linked to the explicit proton transfer reactions along the Cl1 pathway and probed the influence of the redox state of the Mn_4_O_5_Ca cluster by modeling the S_3_Y_z_ and S_3_Y_z_^•^ states.

**Figure 2.**
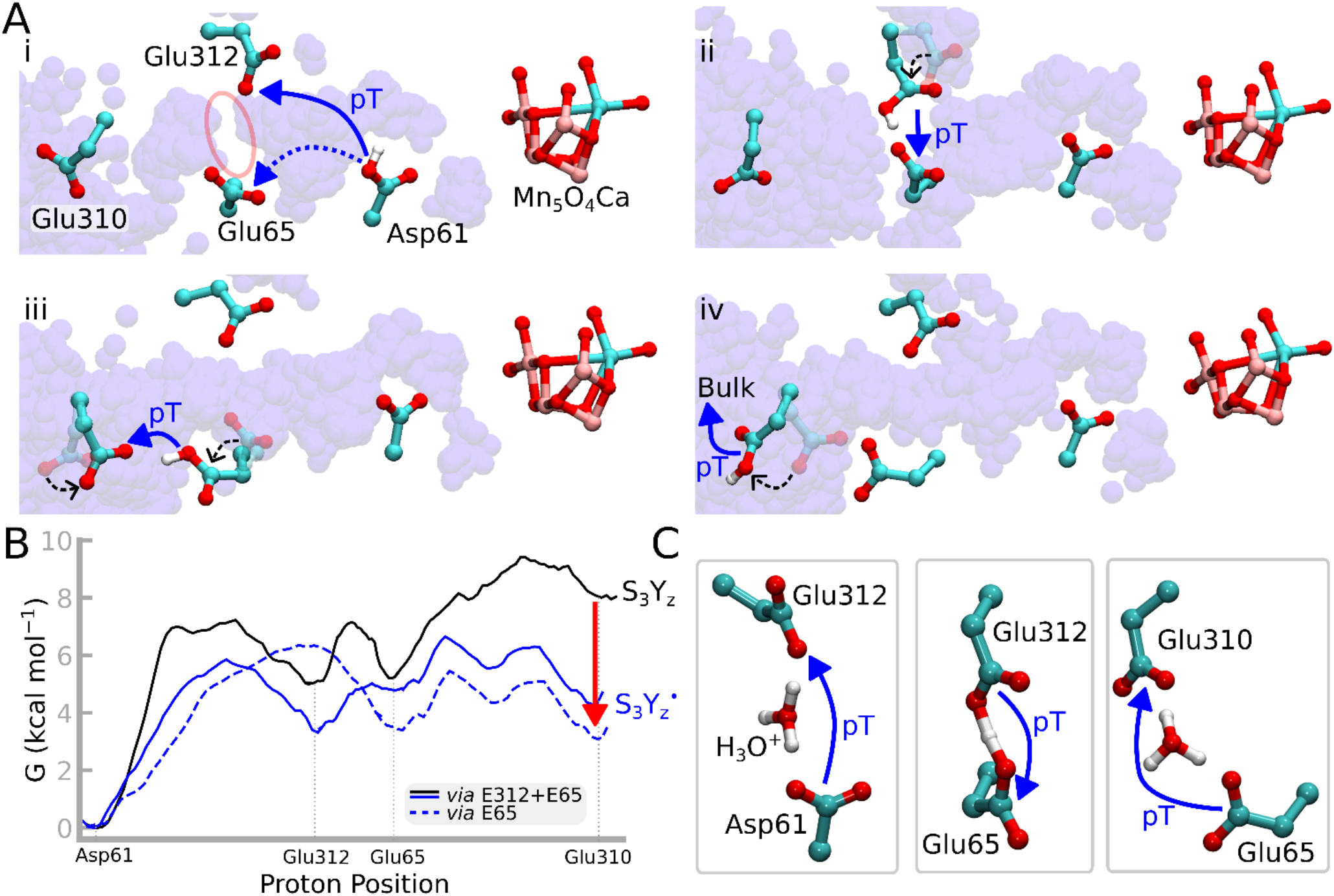
Proton transfer mechanism along the Cl1 pathway. **A)** Proposed proton transfer pathway based on MD simulations. Time averaged overlap of water molecules is shown in transparent blue, indicating the position of the proton transfer channels. The proton channel shows different connectivity depending on the position of Glu65. The area in between Glu312 and Glu65 in particular (highlighted in first panel with a red circle) is critical. Rotations of residues are indicated with dashed black arrows. i) Asp61 donates the proton to either Glu312 or Glu65, ii) Glu312→Glu65, iii) Glu65 rotates, allowing proton transfer to Glu310, and iv) Glu310 donates its proton to the luminal bulk. **B)** Free energy along the proton pathway from Asp61 to Glu310 via Glu312 or Glu65. In the S_3_ state, the proton transfer takes place via Glu312 and Glu65, while in the S_3_^•^ state, the proton can be transferred either via Glu312 and Glu65 or directly via Glu65. **C)** Structure of the transition states based on QM/MM-MD and QM/MM free energy simulations (see also in *SI Appendix*, Fig. S5).

The Y_z_ oxidation involves the transfer of the phenolic proton on Y_z_ to His190_D1_ forming (HisH^+^) (8, 50, 51) and results in an increased proton affinity of Asp61_D1_ (23). This long-range effect of tyrosine oxidation was suggested to arise from a directed electric field that favors opening of the Lys317_D2_/Asp61_D1_ ion-pair and lowers the proton transfer barrier from W3 to Asp61_D1_ (23), [cf. also (31)].

In our classical MD simulations, we note that the protonated Asp61_D1_ forms a tight hydrogen-bonded contact with both Glu65_D1_ (*d*_Asp61-Glu65_ =6.3 Å) and Glu312_D2_ (*d*_Asp61-Glu312_ =6.8 Å) in the S_3_Y_z_^•^ state (*SI Appendix*, Fig. S2A), while Glu65_D1_ points away from Asp61_D1_ in the S_3_Y_z_ state (*SI Appendix*, Fig. S2D). Moreover, Asp61_D1_ and Glu312_D2_ remain around 6-7 Å from each other during all MD simulations, while the Asp61_D1_-Glu65_D1_ distance strongly fluctuates depending on the modeled redox state and the conformation of the Asp61_D1_-Lys317_D2_ ion-pair (*SI Appendix*, Fig. S2D). We note that the modeled S state strongly affects the overall hydration level of the Cl1 channel, with S_3_Y_z_ showing a generally higher hydration level, whilst the oxidation of Y_z_ establishes a strong water-mediated contact between the carboxylate residues (*SI Appendix*, Fig. S4). Protonation of Glu310_D2_/Asp224_PsbO_, which are at the exit of the Cl1 water channel, also leads to an increased hydration within the channel (*SI Appendix*, Figs. S2F and S4).

Our QM/MM free energy simulations show that the water-mediated proton transfer from Asp61_D1_ to Glu312_D2_ has a barrier of *ca*. 7.2 kcal mol^-1^ in the S_3_Y_z_ state, whilst the oxidation of Y_z_ to S_3_Y_z_^•^ and the connected conformational changes in the surrounding ion-pairs, lowers this barrier to *ca*. 5.7 kcal mol^-1^. The water-mediated proton transfer between Asp61_D1_ and Glu65_D1_ also has a modest free energy barrier of *ca*. 6.3 kcal mol^-1^ in S_3_Y_z_^•^, suggesting that it could compete kinetically with the Asp61_D1_ → Glu312_D2_ proton transfer reaction (*SI Appendix*, Fig. S5). In stark contrast, Glu65_D1_ points away from Asp61_D1_ in the S_3_Y_z_ state, which blocks the proton transfer reaction between the residues in this state (Figure 3A, B, *SI Appendix*, Fig. S3 and Fig. S8A). These findings thus suggest that the protons are transferred between Asp61_D1_ and Glu312_D2_/Glu65_D1_ on a nanosecond timescale based on transition state theory (TST) in a process that is favored by the Y_z_ oxidation.

**Figure 3.**
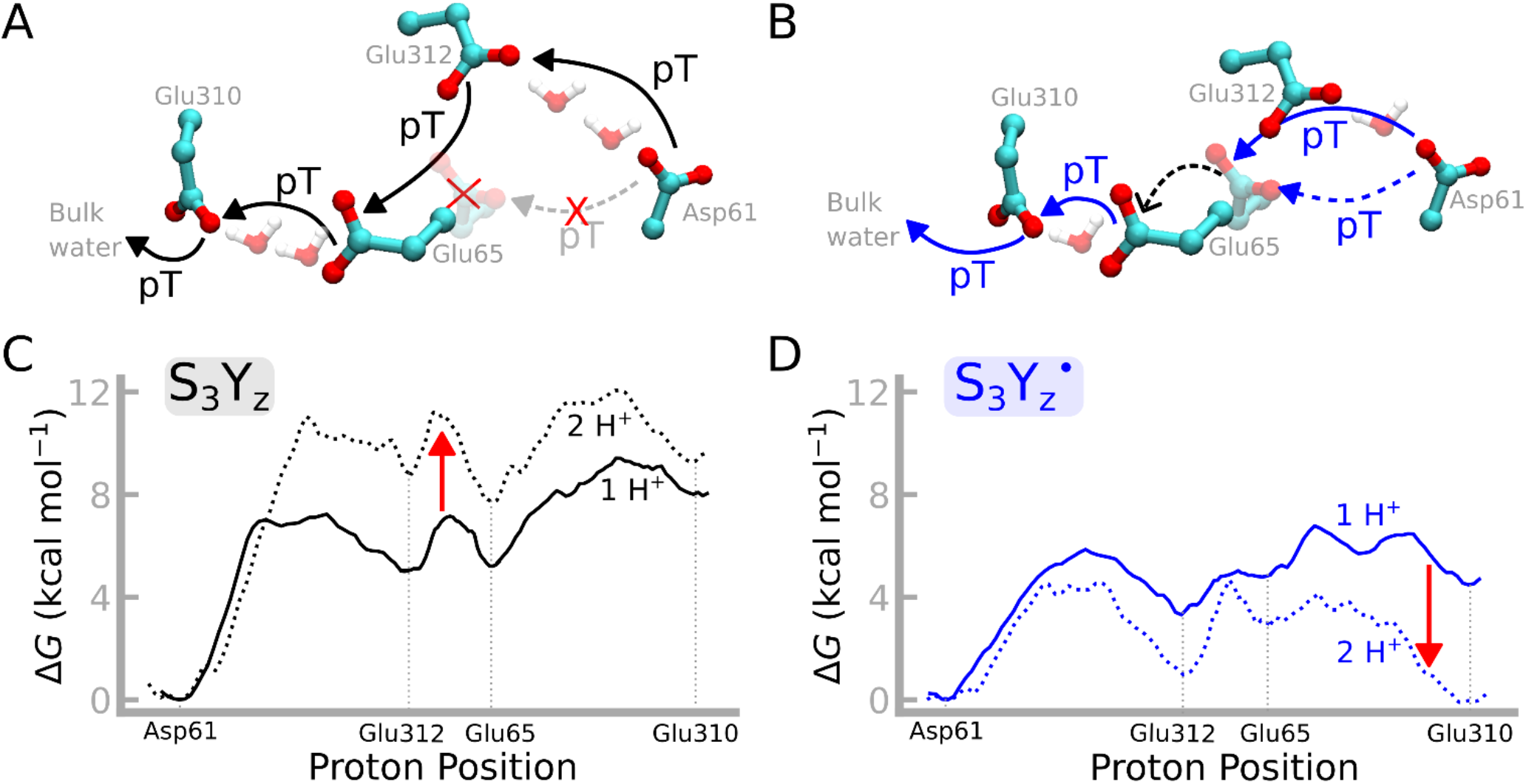
Effect of the S state on the proton pathways. **A-B)** Structural difference between (**A**) the S_3_Y_z_ state and (**B**) the S_3_Y_z_^•^ state. In S_3_Y_z_, Glu65 takes an outward rotated conformation, which renders direct proton transfer from Asp61 to Glu65 unlikely. The opening of Glu65 also drastically changes the hydration within the Cl1 channel (cf. also *SI Appendix*, Fig. S4). In the S_3_Y_z_ state, Asp61→Glu312 and Glu65→Glu312 are connected via two water molecules, while S_3_Y_z_^•^ shows one water molecule between the residues. The figure highlights the four central carboxylates (Asp61, Glu312, Glu65, Glu310) for visual clarity (see *SI Appendix*, Fig. S3 and S4 for further details). **C-D)** Comparison of free energy profiles for **C**) S_3_Y_z_ (in black) and **D**) S_3_Y_z_^•^(in blue). Effect of two protons in the Cl1 channel on the proton transfer energetics. The S_3_Y_z_ state favors one proton in the Cl1 channel, while the S_3_Y_z_^•^ state favors two protons in the Cl1 channel, with the second proton modeled at the Glu312 site.

Our *ab initio* QM/MM-MD simulations capture this spontaneous proton transfer between Glu312_D2_ and Glu65_D1_ on picosecond timescales (*SI Appendix*, Fig. S6C). The proton transfer reaction is consistent with a free energy barrier of 2 kcal mol^-1^ (based on QM/MM-US; *SI Appendix*, Fig. S5B), with similar barriers in both the S_3_Y_z_ and S_3_Y_z_^•^ states.

Remarkably, the protonation of Glu65_D1_ leads to a drastic conformational change of the residue, where its sidechain swings towards Glu310_D2_ (“open gate” conformation; *SI Appendix*, Fig. S2C) and establishes proton connectivity to the luminal bulk (Figure 2A, see also SI Appendix, Fig. S7). During the process, Glu65_D1_ forms an ion-pair with Arg152_PsbO_. In contrast, in the “closed gate” conformation, Glu65_D1_ forms a tight contact with Glu312_D2_, Arg334_D1_, and Asn335_D1_ (see *SI Appendix*, Fig. S8F-H), which blocks the water chain to the bulk, consistent with recent XFEL studies where a rotation of Glu65_D1_ was also observed (22) [cf. also ref. (20)]. This behavior strongly depends on the redox state of the Mn_4_O_5_Ca cluster, with the Y_z_ oxidation favoring the protonation of Glu65_D1_ (see above). We propose that this protonation state dependent modulation of the channel hydration could function as a ratchet to prevent the back-reaction of proton transfer from the luminal side to the oxygen-evolving center.

We find that the rotation of Glu65_D1_ in the S_3_Y_z_^•^ state also affects the proton transfer barrier to Glu310_D2_. In this regard, we obtain a nearly barrierless proton transfer reaction (< 2 kcal mol^-1^) between Glu65_D1_ to Glu310_D2_ in the S_3_Y_z_^•^ state, whilst the barrier is around 4 kcal mol^-1^ in the conformations sampled based on the S_3_Y_z_ state (*SI Appendix*, Fig. S5C). Consistently, we observe a spontaneous proton transfer between Glu65_D1_ and Glu310_D2_ on a *ca*. 35 ps timescale in *ab initio* QM/MM dynamics simulations (QM/MM-MD) in S_3_Y_z_^•^, while the proton remains bound to Glu65_D1_ in S_3_Y_z_ (*SI Appendix*, Fig, S6D). We note that position of Arg115_PsbO_ changes in the simulations of the two S states, with the arginine approaching Glu310_D2_ in S_3_Y_z_^•^ but not in S_3_Y_z_ (*SI Appendix*, Fig. S6B). This redox-triggered arginine flip could thus tune the barrier and driving forces for the proton transfer between Glu65_D1_ and Glu310_D2_ (see below).

The conformation of Glu310_D2_ depends on its protonation state (GluH / Glu^-^), as well as on the protonation state of other nearby titratable residues within the Cl1 channel, suggesting a possible allosteric conformational crosstalk between Glu310_D2_, Glu65_D1_, and Asp61_D1_. When Asp61_D1_ is modeled in a protonated state, Glu310_D2_ is orientated towards the luminal bulk, whilst protonation of Glu65_D1_/Glu312_D2_ favors the “inward conformation” of Glu310_D2_, where it points towards the Mn_4_O_5_Ca cluster (*SI Appendix*, Fig. S8C, D for QM/MM-MD and SI Appendix, Fig. S2B for MD). Moreover, upon protonation of Glu310_D2_, the residue swings towards the luminal bulk (*SI Appendix*, Fig. S8E), whilst its deprotonation favors its inward orientation, a behavior that strongly depends on the conformation of the nearby Asp224_PsbO_ -Arg184_PsbO_ ion-pair. In addition, the “outward conformation” of Glu310_D2_ is favored when the Asp224_PsbO_ -Arg184_PsbO_ ion-pair is in a closed conformation (see *SI Appendix*, Fig. S8I,J). Our simulations suggest that from Glu310_D2_, the proton is transferred either directly or via the conserved Asp224_PsbO_ to the bulk water. In the S_3_Y_z_ state we observe spontaneous rapid proton transfer from Glu310_D2_ to Asp224_PsbO_ (within <20 ps), while the S_3_Y_z_^•^ state supports the formation of a stable Zundel ion (H_5_O_2+_) that can be further transferred to the luminal bulk, with low free energy barriers of only 2-5 kcal mol^-1^. Figure 4A shows an overview of the proposed cluster of carboxylates involved in the proton transfer reaction from the Mn_4_O_5_Ca cluster to the luminal bulk [cf. also Figure 2S, and *SI Appendix*, Fig. S7, S5].

**Figure 4.**
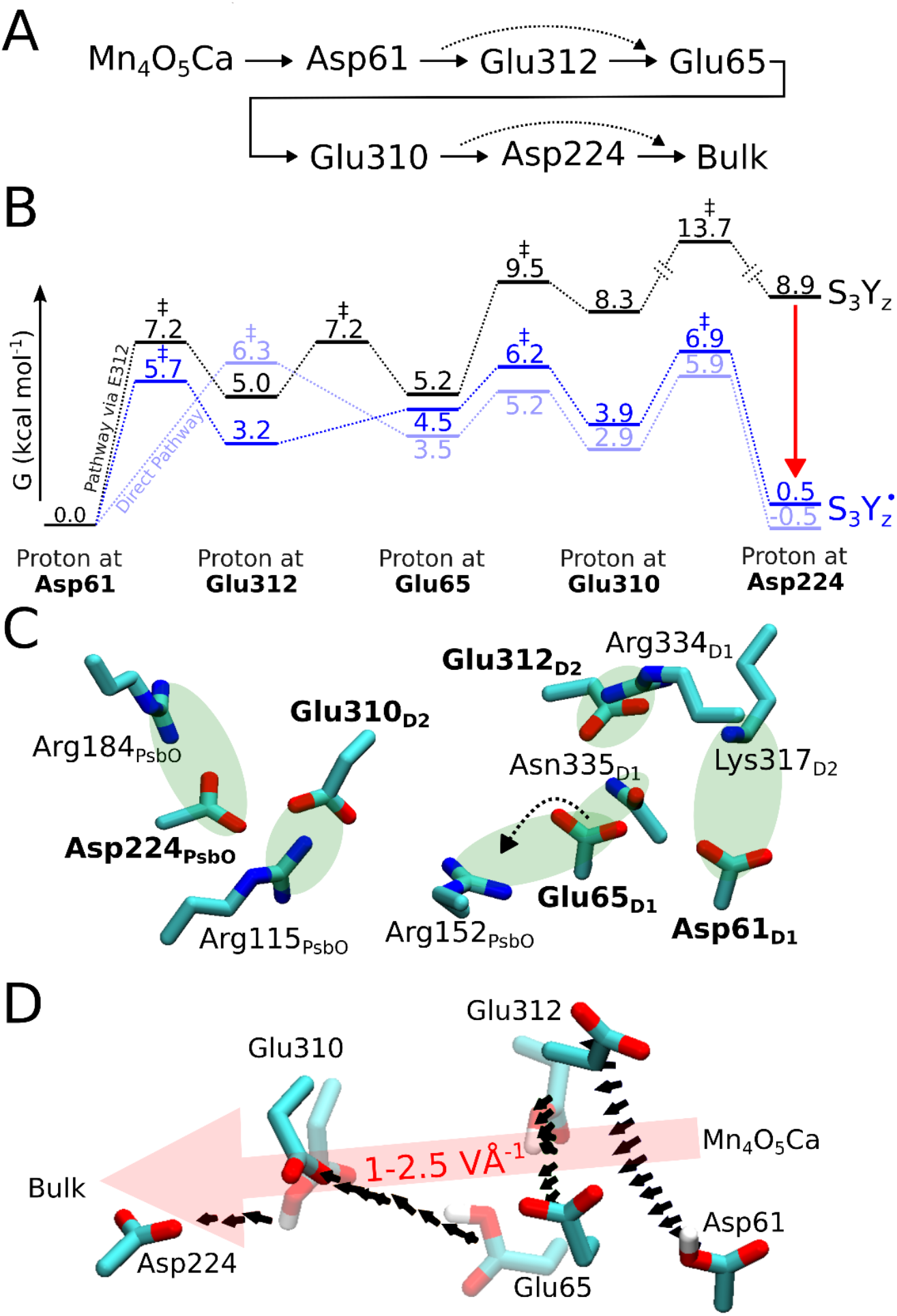
Overview of central residues, free energy profiles, and electric field effects during proton transfer along Cl1 pathway. **A)** Proposed proton transfer pathway from the Mn_4_O_5_Ca cluster to the luminal bulk. **B)** Free energy profile of proton transfer from Asp61 to Glu310 in the S_3_Y_z_ state (in black) and S_3_Y_z_^•^ state (in blue) (see also Figure 2B and *SI Appendix*, Fig. S5 for further details). In comparison to Figure 2B this energy diagram is extended to include Asp224. **C)** Ion-pair contacts (in green) within the central carboxylate cluster. Asp61 forms an ion-pair with Lys317, Glu312 with Arg334, Glu65 with Asn335 (closed conformation) and with Arg152 (open conformation), while Glu310 interacts with Arg115 and Asp224 with Arg184. **D)** Electric fields along the proposed proton pathway (see also *SI Appendix*, Fig. S12 for QM/MM-MD and *SI Appendix*, Fig. S11 for classical MD).

As discussed above, the dynamics and proton transfer barriers are modulated by the conformation of a cluster of positively charged residues (Lys317_D2_, Arg334_D1_, Arg115_PsbO_, Arg152_PsbO_, Arg184_PsbO_) that form ion-pairs with the carboxylates along the Cl1 channel (Figure 4C). We note that Lys317_D2_ forms an ion-pair with Asp61_D1_ and undergoes conformational changes to the open state upon Y_z_ oxidation (*SI Appendix*, Fig. S2K), consistent with our previous findings (23). Moreover, in the deprotonated form, Glu312_D2_ establishes tight ion-pair with both Arg334_D1_ and Lys317_D2_ (*SI Appendix*, Fig. S2H,I), whilst upon protonation, Arg334_D1_ swings towards Glu65_D2_. Similarly, the arginines, Arg115_PsbO_ and Arg152_PsbO_, are located in loop regions that allow the residues to undergo major conformational changes in our atomistic MD simulations. To this end, the two arginines establish ion-pairs with Glu65_D1_ or Glu310_D2_, modulating their dynamics as well as their proton affinities. We note that the described charge elements are highly conserved throughout different organisms, with the exception of Arg115_PsbO_, which is replaced by a glutamate in some organisms (see *SI Appendix*, Fig. S14).

Finally, we also explored the dynamics of the proton transfer reaction, when two protons occupy the Cl1 channel simultaneously, *i*.*e*., by protonating Asp61_D1_/Glu312; Asp61_D1_/Glu65_D1_; or Glu312/Glu65_D1_ at the same time. In such conditions, we observe an overall lower hydration level and disassembly of the water chains connecting the residues. In the QM/MM-MD simulations, the proton from Glu65_D1_ moves spontaneously to Asp61_D1_, or leads to conformational changes around Glu312_D2_, where the residue swings towards the nearby Cl^-^ ion (*SI Appendix*, Fig. S9D). We note that this conformational change together with the altered electrostatic interaction decreases the barrier for nearly all proton transfer steps by a few kcal mol^-1^ (Figure 3C,D and *SI Appendix*, Fig. S9), whilst the barrier for proton transfer between Glu312_D2_ and Glu65_D1_ increases in the S_3_Y_z_^•^ state relative to S_3_Y_z_. These findings suggest that Glu312_D2_ could function as a potential proton storage site, but also that that the Cl1 channel may store and eject only one proton at the time (see *SI Appendix*, Fig. S10).

Taken together, the free energy landscape determined here suggests that the proton transfer from Asp61_D1_ to Glu310_D2_ has a barrier of around 6 kcal mol^-1^ and a Δ*G* of *ca*. 3 kcal mol^-1^ (Figure 2B, Figure 4B) in the S_3_Y_z_^•^ state, which could lead to an effective proton release on the ns-μs timescale, following the oxidation of Y_z_, while further proton transfer to Asp224 favors the reaction even more (Figure 4B). However, our simulations also captured large conformational dynamics linked to the reaction, which may influence the effective barrier transitions by entropic effects. In contrast, the proton transfer in the S_3_Y_z_ state showed a larger overall barrier of around 10 kcal mol^-1^, comparable to a μs-ms timescale process, while the overall reaction is highly endergonic, which thermodynamically disfavors the reactions (Figure 2B, Figure 4B).

Experimentally the proton release from the Mn_4_O_5_Ca cluster takes place on 30-200 μs, depending on the S state (10), with enthalpic barriers of around 7 kcal mol^-1^, and also rather large entropic contributions to the activation energy (*T*Δ*S* ∼6.5 kcal mol^-1^) (31, 52) that could arise from the conformational switching dynamics described above.

### Electric field effects are the driving force for the proton transfer reactions

To probe the physical basis for the proton transfer reactions and its redox-state dependence, we analyzed electric fields along the Cl1 channel, and studied how the barrier of the proton transfer reaction is affected by conformational changes in the ion-pairs.

We find that the electric field points along the proposed proton pathway in the Cl1 channel, with absolute field strengths of 1-2 V Å^-1^, while the oxidation of the Y_z_ (Tyr_z_^•^/HisH^+^) strongly enhances the field effects by *ca*. 0.5 V Å^-1^ (Figure 4B; *SI Appendix*, Fig. S11B). We observe similar electric field effects in both our classical MD simulations (*SI Appendix*, Fig. S11), as well as in the QM/MM-MD simulations (Figure 4, *SI Appendix*, Fig. S12B-E), suggesting that the overall trends are robust.

More specifically, our calculations suggest that the redox-triggered opening of the Lys317_D2_/Asp61_D1_ ion-pair enhances the electric field along the W3→Asp61_D1_ pathway [see also (23)], but also along the Asp61_D1_→Glu312_D2_→Glu65_D1_→Glu310_D2_ pathway. These field effects are likely to result in conformational changes of the ion-pairs within the channel, and in turn could modulate the proton transfer barrier via electrostatic tuning effects. We observe similar electric field effects in both the S_2_→S_3_ and S_3_→S_4_ transitions, suggesting that the proton release step could follow similar principles also in other steps of the photocycle.

To understand how the ion-pair conformations modulate the proton transfer barriers, we introduced perturbations along the Asp61_D1_-Lys317_D2_ distance, by gradually opening the ion-pair and computing the response on the thermodynamic driving force (Δ*E*) and reaction barrier (Δ*E*^‡^) for the proton transfer reaction. The Δ*E vs*. Δ*E*^‡^ relationship provides a basis for understanding rate laws underlying the proton transfer reactions. Thus, the proton transfer barrier between Asp61_D1_ and both Glu312_D2_/Glu65_D1_ show an inverse linear dependence on the ion-pair Asp61_D1_-Lys317_D2_ (1/*d*), with a Brønsted slope of *β=*1.4/0.6 (ΔΔ*E*^‡^ *= β*ΔΔ*E* + *α*). Moreover, the Asp61_D1_ → Glu312_D2_ and Glu65_D1_ → Glu310_D2_ proton transfer barriers show an inverse parabolic dependence on 1/*d*. This observation is consistent with an electric field arising between point charge interactions at close distances and charge-dipole interactions further away, respectively (*SI Appendix*, Fig. S13). The absolute shifts in barriers and driving forces are rather large as the introduced structural perturbations lead to large shifts in electronic energies (see *Methods*), but nevertheless result in linear or parabolic relationships, as expected based on the electrostatic tuning principles.

These observations are similar to our previous findings for other systems (53), which suggest that directed electric field effects favor the formation of water arrays between proton donors and acceptors, and lower the barrier of water-mediated proton transfer reactions.

## Discussion

We have studied here the proton release mechanisms in PSII by combining QM/MM simulations with atomistic molecular dynamics simulations and analysis of electric fields. In this regard, we showed that the light-triggered oxidation of Y_z_ establishes an electric field that favors proton transfer from the Mn_4_O_5_Ca center *via* Asp61_D1_ to Glu65_D1_, which acts as a local molecular gate in the Cl1 channel.

Protonation of Glu65_D1_ leads to a conformational change that establishes a connectivity to the luminal bulk. Although hydrogen positions (and thus protonation states) are not currently resolved in the x-ray structures of PSII (but cf. Ref. (54) released upon finalizing this work), our findings independently support key conformational changes observed in the XFEL structures. Our proposed pathways are indirectly supported by several previous mutagenesis and spectroscopic studies (39, 45, 49, 55-57). In this regard, the D61A substitution decreases the activity of PSII and drastically slows down the oxygen release (39, 45, 49, 55, 56), while mutation of both Glu65_D1_ and Glu312_D2_ also significantly perturb the water oxidation reaction (45, 56, 58), especially the last steps of the photocycle. These findings are also in line with our previous work, suggesting that both Asp61_D1_ and Glu65_D1_ are involved in the water oxidation reaction (23). Interestingly, mutation of Glu312_D2_ appear less detrimental than those of Glu65_D1_ (56), indicating that Glu65_D1_ could establish the primary proton pathway, Glu312_D2_ may serve as a potential proton storage site, as suggested in our current work.

The Glu65_D1_/Glu312_D2_ site was recently suggested to undergo stepwise deprotonation and re-protonation reactions (31). However, the interpretation of the IR spectra was recently challenged (32), with an alternative suggestion that the spectral shifts result from conformational changes that could regulate the proton transfer. Our current work strongly supports that the Glu65_D1_/Glu312_D2_ sites undergo both protonation and conformational changes that are likely to contribute to the observed vibrational shifts in the 1707 cm^-1^ range.

The conformational changes of Glu65_D1_ as predicted by our simulations are also observed in recent XFEL structures of the S_3_Y_z_ → S_3_Y_z_^•^ transition (see *SI Appendix*, Fig. S15). We note that our simulations predict somewhat larger conformational changes than those observed experimentally, which could result from averaging effects, and accumulation of different photocycle states during high S states.

All simulations performed here are based on the dark state x-ray structure, and thus not biased by the conformational changes resolved in the XFEL structures. We thus argue that our simulations provide important independent validation to these recent experiments, which is valuable given the technical challenges in data refinement and accumulation of different S states in the XFEL experiments. Our simulations, although limited to the μs timescales, probe the effects of redox and protonation states, and their underlying physical principles, whilst our QM/MM simulations reveal the energetics of the proton transfer process.

We note that the dynamic water networks in combination with re-arrangement of ion-pairs resolved here indicate that large entropic effects are involved in the proton transfer reactions, consistent with the rather large experimental *T*Δ*S* (∼7 kcal mol^-1^) contributions observed for the S_3_Y_z_ → S_4_Y_z_ transition (31).

Taken together, our findings suggest that the complex allosteric crosstalk within the Cl1 channel is controlled by the redox state that establishes an electric field, which in turn controls the proton transfer reactions and the conformation of ion-pairs. We suggest that the functional elements identified here could provide a kinetic gate that prevents the back-transfer of protons from the luminal bulk to the OEC, particularly under high *pmf* conditions. Our findings thus provide a new physical understanding of the underlying forces that result in these changes and show that the Cl1 pathway supports proton transfer from the substrate water molecules to the luminal bulk.

## Conclusion

We showed here that Photosystem II employs the Cl1 channel to catalyze the stepwise light-driven deprotonation of substrate water molecules during the photocycle. The proton transfer is tightly controlled by redox-gates that undergo hydration transitions and conformational changes in ion-pairs, both caused by electric fields arising from the redox state of the Mn_4_O_5_Ca center and oxidation of Y_z_. We suggest that the Glu65_D1_ gate identified here modulates the barrier for proton transfer along the Cl1 channel and connectivity to the luminal bulk. This could be important for regulating the back-transfer of protons under physiological conditions, whilst Glu312_D2_ could act as a potential proton storage site. The elucidated principles suggest that several bioenergetic systems might employ electric field effects, conformational changes of ion-pairs, and hydration transitions to control proton transfer reactions (23, 53, 59, 60).

## Materials and Methods

### Molecular dynamics simulations

The x-ray structure of the PSII from *T. vulcanus* (PDB ID: 3WU2) (12) was embedded in a 1-palmitoyl-2-oleoyl-*sn*-glycero-3-phosphocholine (POPC) membrane, and solvated with TIP3P water molecules and 100 mM NaCl. The total system comprised around 535,000 atoms (see *SI Appendix*, Fig. S14 for a sequence analysis and *SI Appendix*, Fig. S15 for a structural comparison). The cofactors were modeled using in-house force field parameters (23, 61), while the remaining system was modeled using the CHARMM36m force field (62). MD simulations were performed in each of the following states, S_2_Y_z_^•^, S_3_Y_z_, S_3_Y_z_^•^, S_4_Y_z_, as well as different protonation states (Asp61_D1_, Glu312_D2_, Glu65_D1_, Glu310_D2_, and Asp224_PsbO_), thus modeling the stepwise proton transfer along the Cl1 pathway. In total, 48 independent MD simulations were performed. Each simulation was 200 ns, yielding in total *ca*. 10 μs. All simulations were performed using NAMD2.14 and NAMD3.0alpha9 (63, 64), and analyzed using VMD (65). The simulations were carried out in an *NPT* ensemble at *T*=310 K and *p*=1 atm. The particle mesh Ewald (PME) method with a 1 Å grid separation was employed to account for long-range electrostatic interactions. Prior to the 200 ns production runs, the MD system was relaxed for 4 ns with harmonic restraints on the protein of 1 kcal mol^−1^ Å^−1^, followed by 30 ns of equilibration without restraints. All classical MD simulations are reported in *SI Appendix*, Table S1. The MD simulation system is shown in *SI Appendix*, Fig. S1A.

### QM/MM simulations

Hybrid QM/MM calculations were performed using structures obtained from the MD simulations (see above). The QM/MM model comprised ca. 16,500 MM atoms and 258-276 QM atoms, modeling the Cl1 pathway. The QM region included the proton acceptors (Asp61_D1_, Glu312_D2_, Glu65_D1_, Glu310_D2,_ Asp224_PsbO_), residues directly coordinating these (Lys317_D2_, Arg334_D1_, Asn335_D1_, Arg152_PsbO_, Arg115_PsbO_), as well as Asp59_D1_, Arg64_D1_, Tyr107_D1_, Thr107_PsbO_, and Tyr151_PsbO_, together with 21-27 water molecules. The proton transfer between Asp61_D1_ – Glu312_D2_, Glu312_D2_ – Glu65_D1_, Asp61_D1_ – Glu65_D1_, Glu65_D1_ – Glu310_D2_ and Glu310_D2_ – Asp224_PsbO_ were modeled in individual QM/MM simulations. The QM region was described at the B3LYP-D3/def2-SVP level of theory (66-69). Link atoms were added between the C_α_ and C_β_ atoms on the amino acid side chains. For equilibration, the QM/MM system was gradually heated from *T*=0 K to 310 K using a timestep of 0.1 fs over 310 fs, followed by QM/MM dynamics at *T* = 310 K with a 1 fs integration timestep. During the QM/MM simulations, a 10 Å sphere around the QM region was allowed to fully relax, while the remaining protein was kept frozen. The QM/MM calculations were performed using FermiONs++ (70, 71). Details of the QM/MM simulations are listed in *SI Appendix*, Table S2, and the computational models are shown in *SI Appendix*, Fig. S1B,C. The free energy of the proton transfer along the Cl1 pathway was studied using umbrella sampling (US), with the parameters matching those of the unbiased QM/MM simulations. The reaction coordinate was defined as a linear combination of all distances involved in bond formation/breaking along the proton transfer pathway (see *SI Appendix*, Fig. S6A). To this end, the proton transfer reaction was modeled using 6-17 Windows with a force constant of 100 kcal mol^-1^ Å^-2^ and the energy profiles were calculated using the weighted histogram analysis method (WHAM) (72). See *SI Appendix*, Table S3 for details of the free energy calculations. Representative structures (30 snapshots in total) were extracted from simulations of the reactant state, transition state, and product state along the umbrella sampling, to study the influence of the Asp61-Lys317 ion-pair distance on the proton transfer barriers.

## Acknowledgments

This project received support from the Knut and Alice Wallenberg Foundation (V.R.I.K. grant: 2019.0251), the Swedish Research Council (V.R.I.K.), the Royal Society (Royal Society Research Professorship 2024 to AWR) Biotechnology and the Biological Sciences Research Council (BBSRC) (A.W.R. grants: BB/R001383/1 and BB/R00921X) and Leverhulme Trust Grant RPG-2022-203 to AWR. V.R.I.K. also acknowledges the DFGs support within the Mercator Fellow Program to SFB1078. Computational resources were provided by the Leibniz Rechenzentrum (LRZ, project:pr83ro), the National Academic Infrastructure for Supercomputing in Sweden (NAISS 2023/1-31, NAISS 2024/1-28) and the Swedish National Infrastructure for Computing (SNIC2022/1-29).

