## Supplementary material for "Mechanism of Proton Release during Water Oxidation in Photosystem II": SI Appendix

### Supporting Information for
Mechanism of Proton Release during Water Oxidation in Photosystem II

Friederike Allgöwer[1], Maximilian C. Pöverlein[1], A. William Rutherford[2], Ville R. I. Kaila[1,*]

[1] Department of Biochemistry and Biophysics, Stockholm University, 10691 Stockholm, Sweden.
[2] Department of Life Sciences, Imperial College London, London SW7 2AZ, United Kingdom.

*

**This PDF file includes:**

**Figure S1.** Computational models.
**Figure S2.** Analysis of distances from MD simulations.
**Figure S3.** Structural comparison of the $S_3Y_z$ and $S_3Y_z^{\bullet}$ states.
**Figure S4.** Hydration levels of the Cl1 channel from QM/MM-MD simulations.
**Figure S5.** Free energy profiles of proton transfer from QM/MM-US simulations.
**Figure S6.** Definition of reaction coordinate and protonation dynamics.
**Figure S7.** Water connectivity along the Cl1 pathway in the QM/MM-MD simulations.
**Figure S8.** Dynamics of the Glu65 gate from QM/MM-MD simulations.
**Figure S9.** Effect of two protons in the Cl1 channel.
**Figure S10.** Schematic representation of the energy barriers for proton transfer with two protons in the Cl1 channel.
**Figure S11.** Analysis of electric field effects along the Cl1 pathway from MD simulations
**Figure S12.** Analysis of electric field effects along the Cl1 pathway from QM/MM-MD simulations.
**Figure S13.** Reaction barriers as a function of driving forces from perturbation analysis.
**Figure S14.** Multiple sequence alignment.
**Figure S15.** Structural comparison between predicted conformational changes and x-ray structures.

**Table S1.** List of classical MD simulations.
**Table S2.** List of QM/MM-MD simulations.
**Table S3.** List of QM/MM free energy simulations (QM/MM-US).

**SI References**

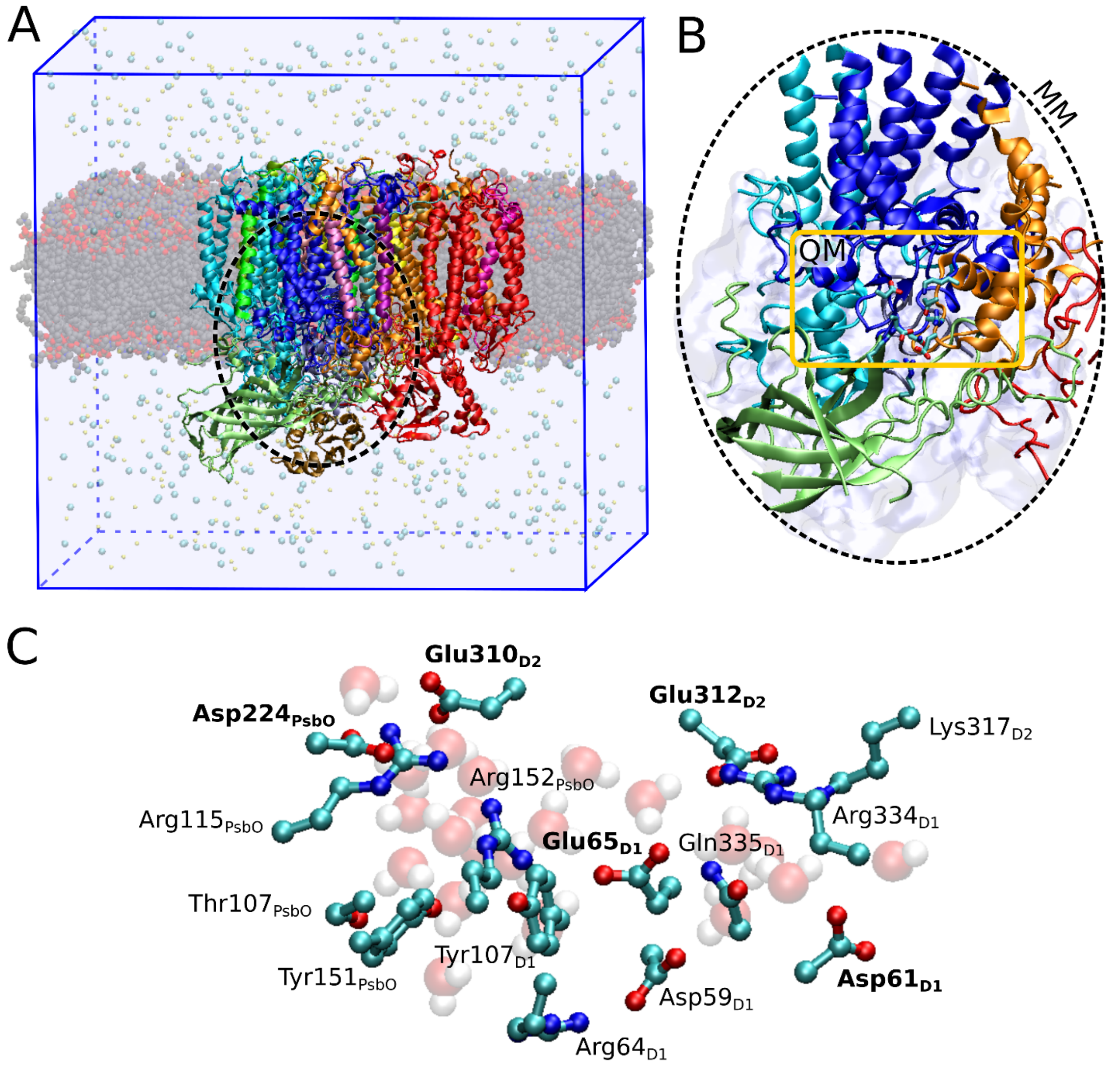


**Fig. S1. Computational models. A)** Atomistic MD model. The protein coordinates (PDB ID: 3WU2(1)) were embedded in a membrane and a water-ion box (see *Materials and Methods*). **B)** The QM/MM model. The MM region was defined as a 23 Å sphere around the QM region (orange box). All atoms around 10 Å from the QM region were allowed to move during the QM/MM simulations. **C)** The QM region of the QM/MM models comprised *ca.* 260 atoms. Residues participating in the proton transfer reactions are labeled in bold.

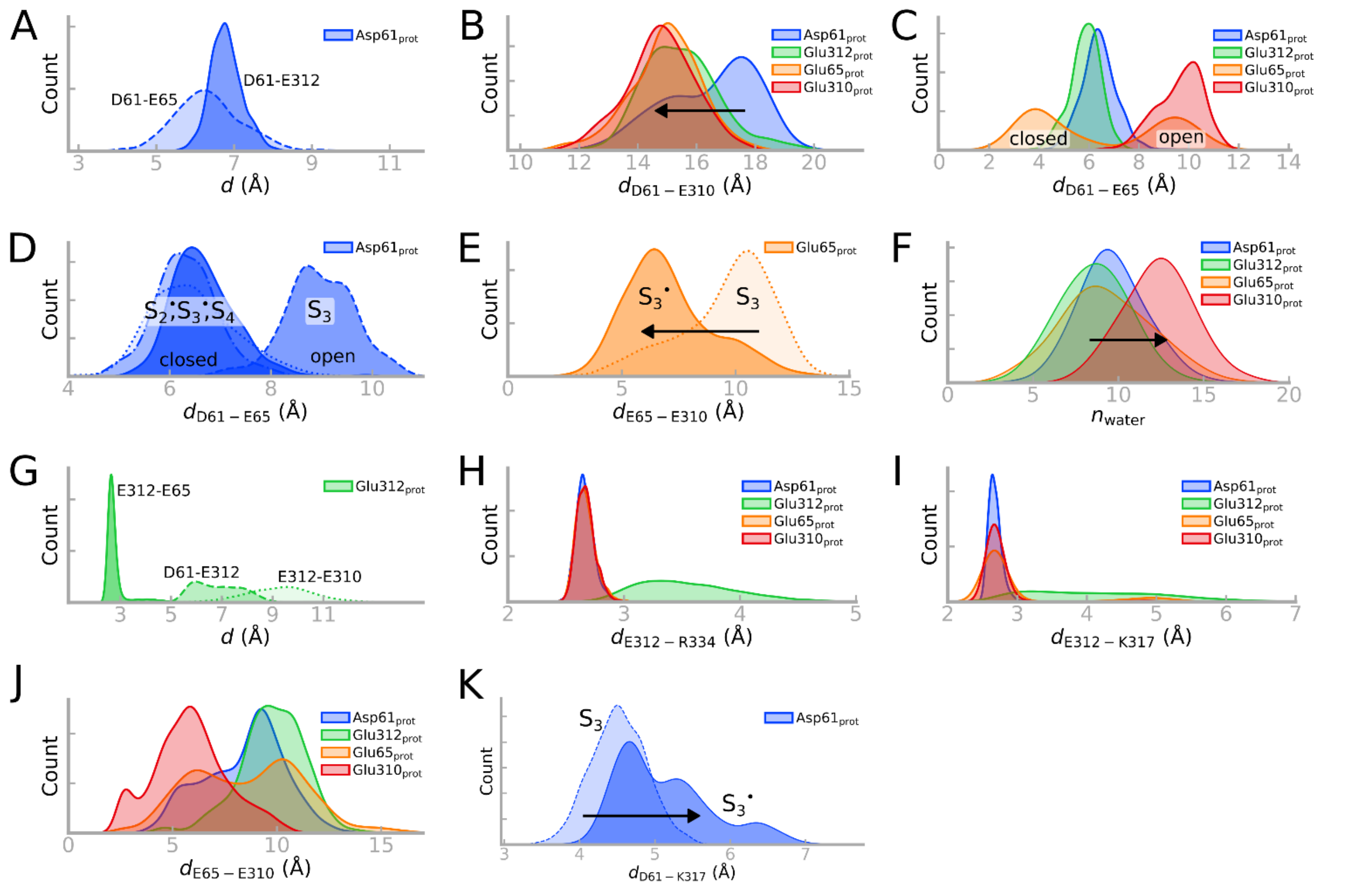


**Fig. S2. Analysis of distances from MD simulations.** The population distributions from MD. Distances were measured between closest heavy atom contacts (OE1, OE2, N). The different colors refer to models with different protonated residues within the Cl1 channel (Asp61 protonated in blue; Glu312 protonated in green; Glu65 protonated in orange; Glu310 protonated in red). **A)** Distances between Asp61-Glu312 (solid) and Asp61-Glu65 (dashed) with protonated Asp61 show similar trends. Therefore, both residues, Glu312 and Glu65, could function as proton acceptors from Asp61. **B)** The Asp61-Glu310 distance shows how Glu310 moves towards Asp61 (and Glu65) after the proton transfer from Asp61. **C)** The distance between Asp61 and Glu65 shows how the Glu65 gate is closed when Asp61 (Asp61$_{prot}$) or Glu312 (Glu312$_{prot}$) is protonated. Upon proton transfer to Glu65 (Glu65$_{prot}$), the residue switches to its open conformation until the protonation of Glu310 (Glu310$_{prot}$). **D)** In the $S_3$ state, Glu65 is in an open conformation, while in other S states the closed conformation is preferred. The Asp61-Glu65 distance is therefore sensitive to the modelled redox state (for Asp61$_{prot}$). **E)** The Glu65-Glu310 distance is shown for $S_3Y_z$ (*avg*. 10.5 Å) and the $S_3Y_z^{\bullet}$ state (*avg*. 6.5 Å). **F)** Average hydration level of the Cl1 channel from MD simulations with different protonation states (Asp61$_{prot,}$ Glu65$_{prot}$, Glu312$_{prot}$, Glu310$_{prot}$). The simulations show averaged data over all S states. The hydration level increases as the proton approaches the luminal bulk (Glu310$_{prot}$, in red). **G**) Protonation of Glu312 favors a short Glu65-Glu312 distance, while the Asp61-Glu312 and Glu312-Glu310 distances are in comparison longer, and indicating that proton transfer is likely to occur via Glu65. **H-I)** Glu312 forms an ion-pair with Arg334 (H) or Lys317 (I), except when the residue is protonated (Glu312$_{prot}$; green). **J)** Protonation of Glu65 or Glu310 favors close contacts between Glu65 and Glu310. **K)** The Asp61-Lys317 ion-pair opens up upon oxidation of $Y_z$.

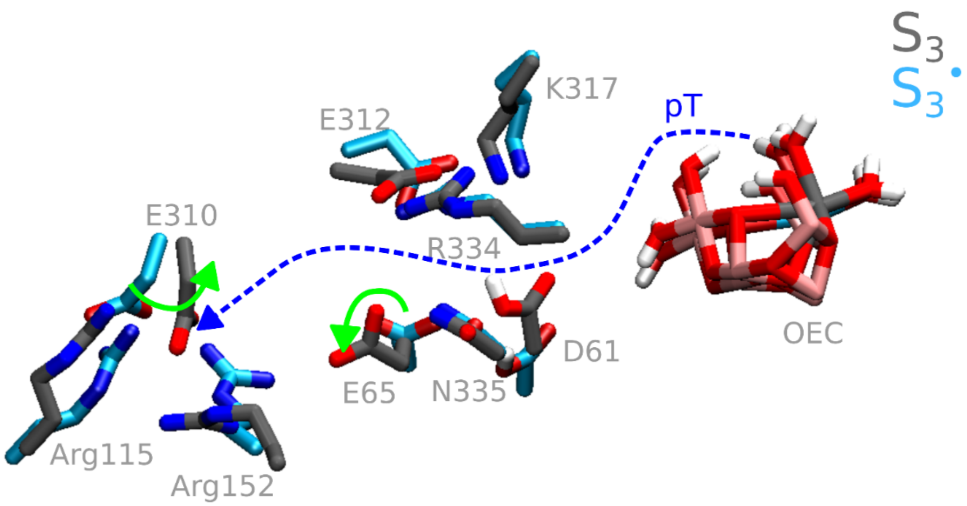


**Fig. S3. Structural comparison of the $S_3Y_z$ and $S_3Y_z^{\bullet}$ state.** Structural overlay of the $S_3Y_z$ state (open Glu65 conformation; in grey) and the $S_3Y_z^{\bullet}$ state (closed Glu65 conformation; in blue). The central carboxylates are shown (Asp61, Glu312, Glu65, Glu310), together with their corresponding ion-paired contacts (Lys317, Asn335, Arg334, Arg152, Arg115). Rotations of the residues are indicated with solid green arrows, and the general proton transfer direction from the $Mn_4O_5Ca$ cluster to the bulk water is shown with a dashed blue arrow.

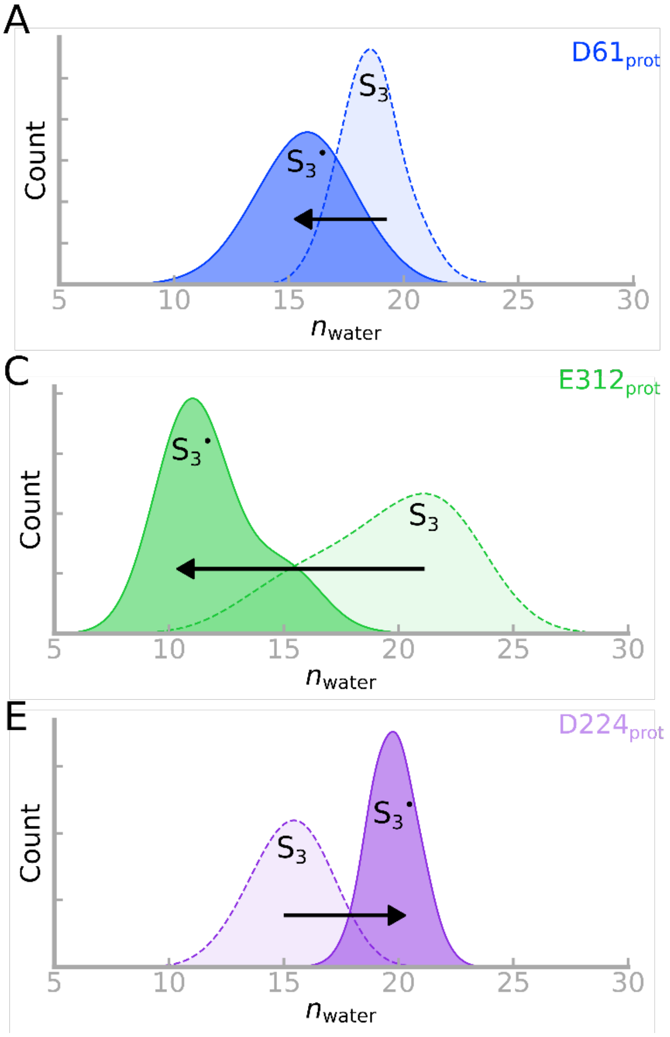


**Fig. S4. Hydration levels of the Cl1 channel from QM/MM-MD simulations.** The protonation states of (**A**) Asp61$_{prot}$, **B)** Glu65$_{prot}$, **C)** Glu312$_{prot}$, **D)** Glu310$_{prot}$ and **E)** Asp224$_{prot}$ are shown. The $S_3Y_Z$ state shows an overall higher hydration level relative to the $S_3Y_Z^{\bullet}$ state, except when the proton is at Asp224 (purple). The hydration level increases also as the proton moves towards the luminal bulk (Glu310$_{prot}$; in red). The QM/MM shows similar trends as observed in the classical MD simulations (Fig. S2F).

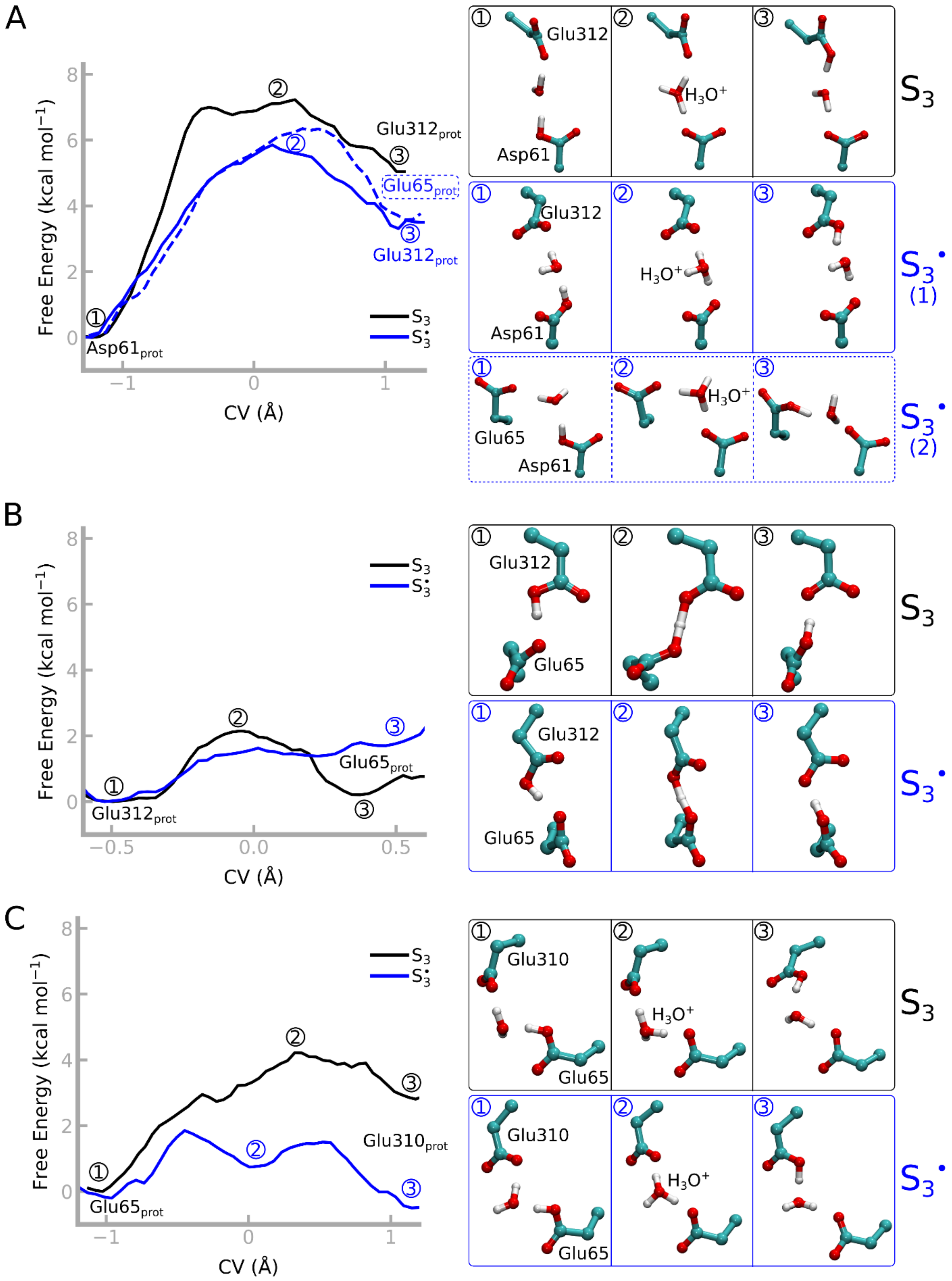


**Fig. S5. Free energy profiles of proton transfer from QM/MM-US simulations**. Proton transfer along **A)** the Asp61→Glu312/Glu65, **B)** the Glu312→Glu65, **C)** the Glu65→Glu310 and **D)** the Glu310→Asp224 pathways. *Inset*: Structures of reaction intermediates extracted along the free energy profile for both $S_3$ (black) and $S_3^{\bullet}$ (blue).

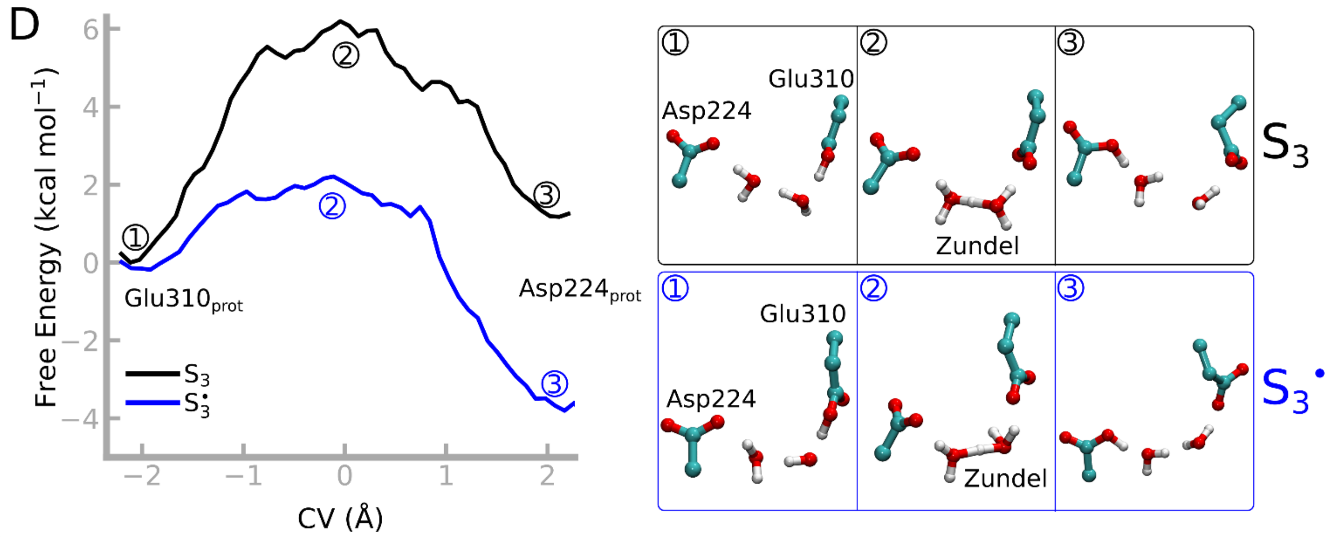


**Fig. S5 *(contd.)*. Free energy profiles of proton transfer from QM/MM-US simulations.** Proton transfer along **A)** the Asp61→Glu312/Glu65**, B)** the Glu312→Glu65, **C)** the Glu65→Glu310 and **D)** the Glu310→Asp224 pathways. *Inset*: Structures of reaction intermediates extracted along the free energy profile for $S_3$ (black) and $S_3^{\bullet}$ (blue).

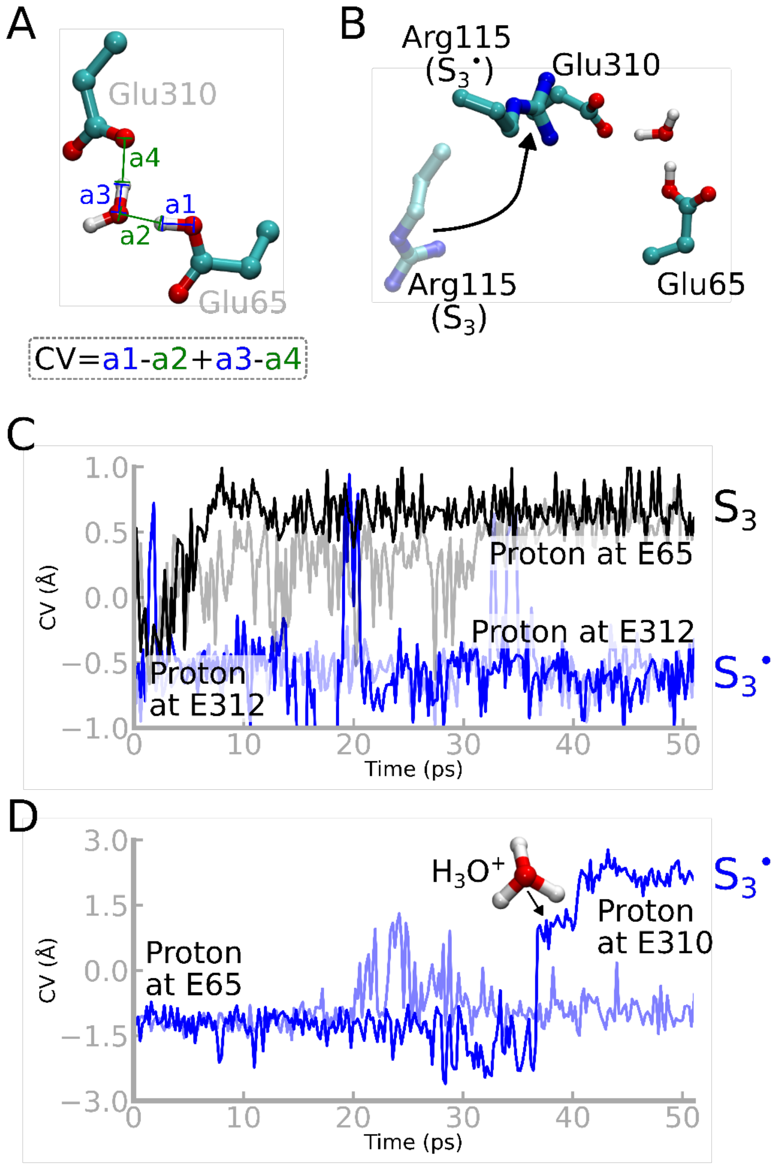


**Fig. S6. Definition of reaction coordinate and protonation dynamics. A)** Definition of the reaction coordinate (collective variable, CV) used in the QM/MM free energy calculations. The proton transfer reaction between Glu65 and Glu310 is used here as an example for the CV definition. Analogous definitions of the CVs were used for all residue pairs within the Cl1 pathway. **B)** In the $S_3Y_z$ state, Arg115 swings outwards, while in the $S_3Y_z^{\bullet}$ state, Arg115 forms a close contact with Glu310. The conformational changes could favor the proton transfer from Glu65 to Glu310 in the $S_3Y_z^{\bullet}$ state. **C)** Proton transfer reactions from QM/MM-MD simulation, initiated with a protonated Glu312. The proton moves from Glu312 to Glu65 in the $S_3Y_z$ state, while the protonated form of Glu312 is favored in the $S_3Y_z^{\bullet}$ state. **D**) Proton transfer reactions from QM/MM-MD simulation, initiated with a protonated Glu65. In one out of the two replicas of the $S_3Y_z^{\bullet}$ state, the proton spontaneously moves to Glu310. Data for the $S_3Y_z$ is not shown, as the proton wire is disassembled (see *main text*).

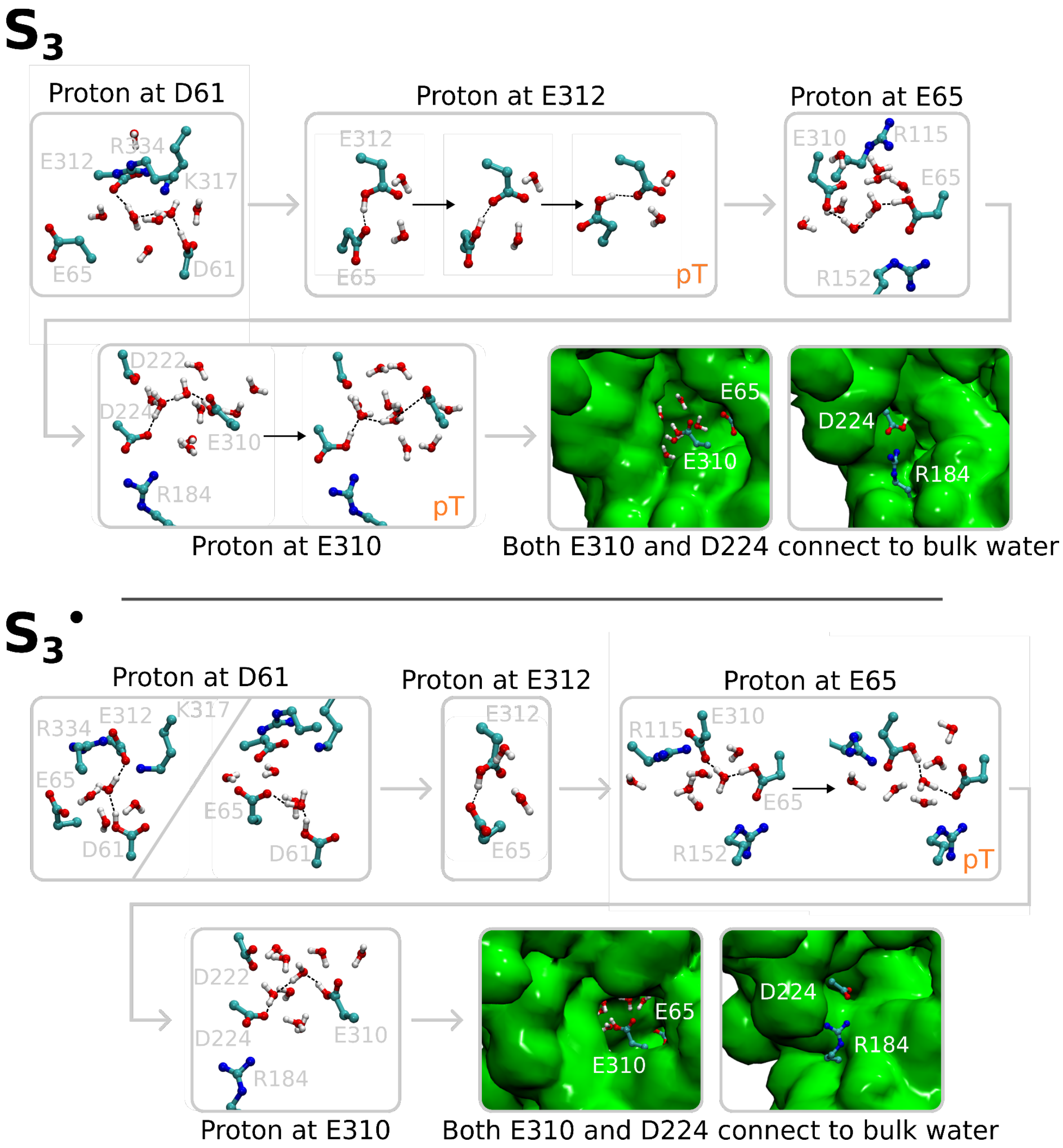


**Fig. S7. Water connectivity along the Cl1 pathway in the QM/MM-MD simulations.** Upon protonation of Asp61, the simulations show a stronger proton wire in the $S_3Y_z^{\bullet}$ state relative to $S_3Y_z$, suggesting that the oxidation of $Y_z$ favors proton transfer along the Cl1 pathway. The $S_3Y_z$ state enables the proton transfer between Glu312 and Glu65, leading to conformational change of the latter, while in the $S_3Y_z^{\bullet}$ state, the proton moves rapidly between Glu312 and Glu65. Protonation of Glu65, favors the formation of a water chain between Glu65 and Glu310. The water array is longer in the $S_3Y_z$ state relative to $S_3Y_z^{\bullet}$ state, enabling a proton transfer to Glu310 (and/or Asp224). Glu310 and Asp224 are both accessible to the luminal bulk.

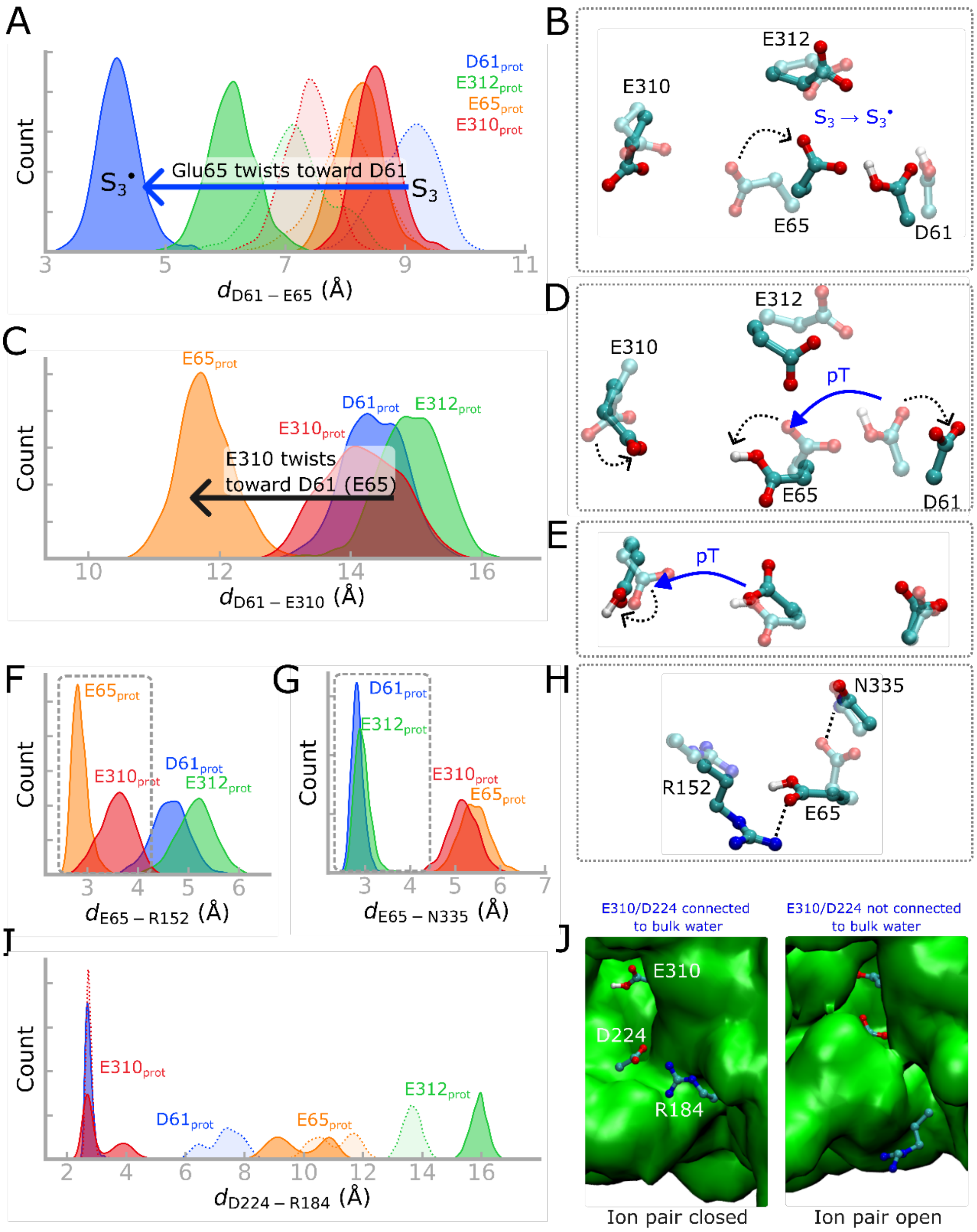


**Fig. S8. Dynamics of the Glu65 gate from QM/MM-MD simulations. A, B)** Comparison of ion-pair distances and representative structures in the $S_3Y_Z$ and $S_3Y_Z^{\bullet}$ states. Protonation of Asp61 shifts the Asp61-Glu65 distance (in blue). The $S_3Y_Z \rightarrow S_3Y_Z^{\bullet}$ transition favors a conformational change around Glu65. The open conformation of Glu65, enables the proton transfer between Asp61 → Glu312 (in $S_3Y_Z$), while the closed conformation enables proton transfer between Asp61 → Glu312 and Asp61 → Glu65 (in $S_3Y_Z^{\bullet}$). **C)** The Asp61-Glu310 distance and **D**) representative structures showing conformational changes of both Glu65 and Glu310 that may facilitate the proton transfer between the residues. **E)** Glu310 swings towards the luminal bulk upon its protonation. **F-H)** Comparison of the "open" and "closed" conformation of Glu65. In the closed conformation, Glu65 coordinates with the nearby Asn335, while in the open conformation, Glu65 coordinates with Arg152. **I-J)** The Asp224-Arg284 ion-pair is located close to the surface of the protein. In the closed conformation of this ion-pair, Glu310 and Asp224 are connected to the luminal bulk, while in the open conformation, the residues remain shielded from the bulk. The colors in panels A-J refer to models with different protonated residues within the Cl1 channel (protonated Asp61 in blue; Glu312 in green; Glu65 in orange; Glu310 in red).

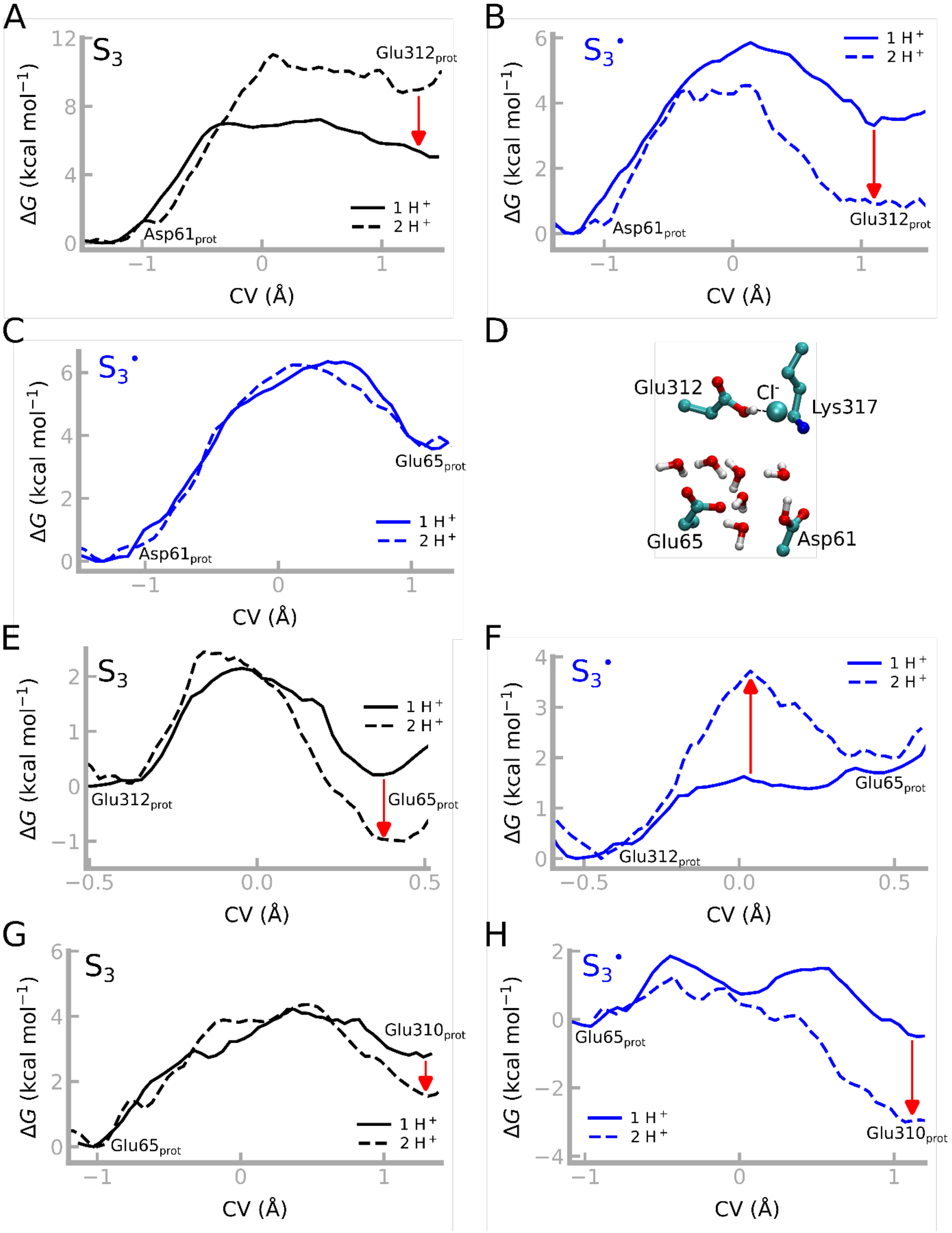


**Fig. S9. Effect of two protons in the Cl1 channel**. The free energy profile for proton transfer from **A-B**) Asp61→Glu312 with Glu65($^{-}$) (*solid*) and Glu65(H) (*dashed*), **C**) Asp61→Glu65 with Glu312($^{-}$) (*solid*) and Glu312(H) (*dashed*), **E-F)** Glu312 → Glu65 with Asp61($^{-}$) (*solid*) and Asp61(H) (*dashed*), **G-H**) Glu65→Glu310 with Glu312($^{-}$) (*solid*) and Glu312(H) (*dashed*), in the $S_3Y_Z$ state (in black), and the $S_3Y_Z^{\bullet}$ state (in blue). An additional proton in the Cl1 pathway, decreases the proton transfer barrier by 2-3 kcal mol$^{-1}$ in most steps. However, the Asp61→Glu65 barrier is unaffected by an additional proton in the channel, with the Glu312→Glu65 barrier increasing by ca. +2 kcal mol$^{-1}$. **D**) QM/MM-MD simulations with protons on both Asp61 and Glu312, leading to a flip of Glu312 towards the $Cl^-$ ion.

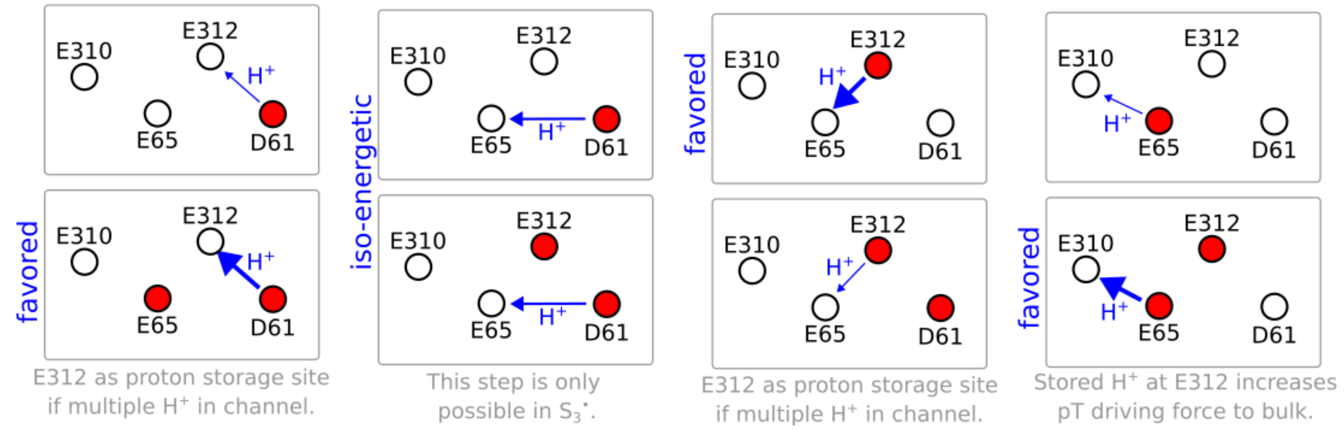


**Fig. S10. Schematic representation of the energy barriers for proton transfer showing the effect of two protons in the Cl1 channel.** The individual energy barriers are shown in Fig. S5 and Fig. S9. The four central carboxylates are shown for visual clarity, protonated carboxylates are marked in red, deprotonated carboxylates in white, proton transfers are indicated by blue arrows. Thick arrows represent highly favored proton transfer reactions, thin arrows less favored reactions. In general, the proton storage at Glu312 is favored when multiple protons reside in the channel.

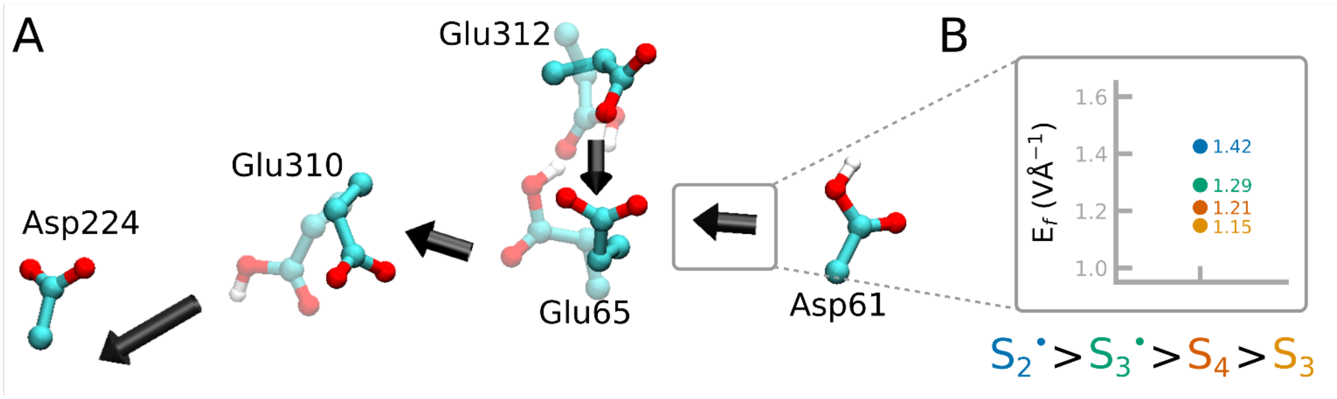


**Fig. S11. Analysis of electric field effects along the Cl1 pathway (from MD simulations).** A) Directionality of electric fields shown at the midpoint between carboxylates along the Cl1 pathway. **B)** Comparison of field strengths depending on the S state shown as an example for a selected probe points (between Asp61-Glu65). A clear trend of $S^{\bullet}$>S is visible, which is consistent also at all other measuring points. Analysis of the electric fields in the QM/MM simulations (Fig. S12) show similar trends.

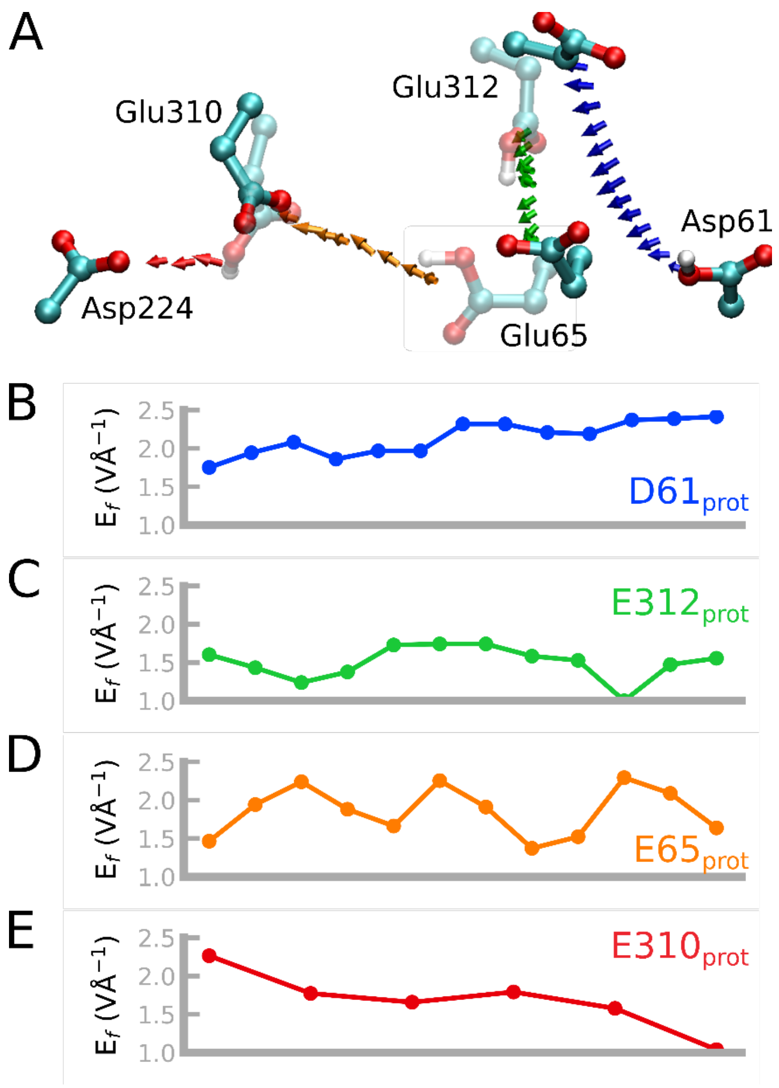


**Fig. S12. Analysis of electric field effects along the Cl1 pathway (from QM/MM-MD simulations). A-E)** Absolute electric fields in the $S_3Y_z^{\bullet}$ state along the Cl1 pathway from QM/MM-MD simulations. The colors indicate the position of the proton: Asp61 protonated in blue; Glu312 protonated in green; Glu65 protonated in orange; and Glu310 protonated in red. Analysis of the electric fields in the MD trajectories (Fig. S11) show similar trends.

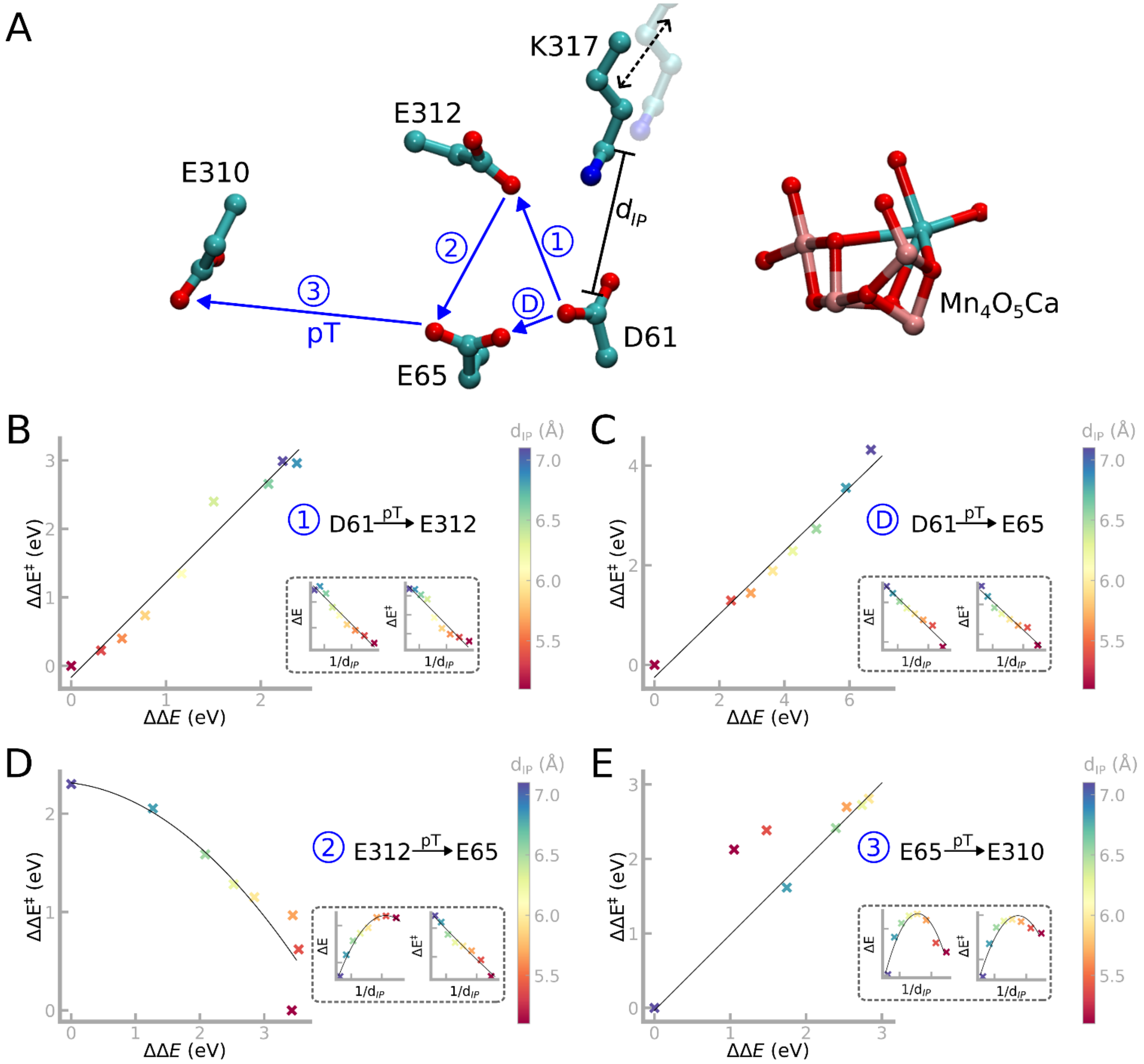


**Fig. S13. Reaction barriers as a function of driving forces from perturbation analysis.** Effect of perturbations in the ion-pair distance (Asp61-Lys317) on the proton transfer energetics. **A)** Labeling of the individual proton transfer steps (1, D, 2, and 3). **B-E**) Barrier as a function of driving force for the proton transfer steps, and their dependence on the Asp61-Lys317 ion-pair distance (*inset*). The ion-pair distance was shifted by displacing the residues sidechains along the Asp61-Lys317 bond vector, followed by estimation of the response on the reaction barrier and driving force by single point calculations. The proton transfer shows a linear barrier dependence on the driving force close to the Asp61/Lys317 ion-pair (Asp61→Glu65 and Asp61→Glu312), and a parabolic dependence on the driving force further along the pathway (Glu312→Glu65). The insets in each panel show the driving force and barrier as a function of each specific ion-pair distance ($d_{IP}$).

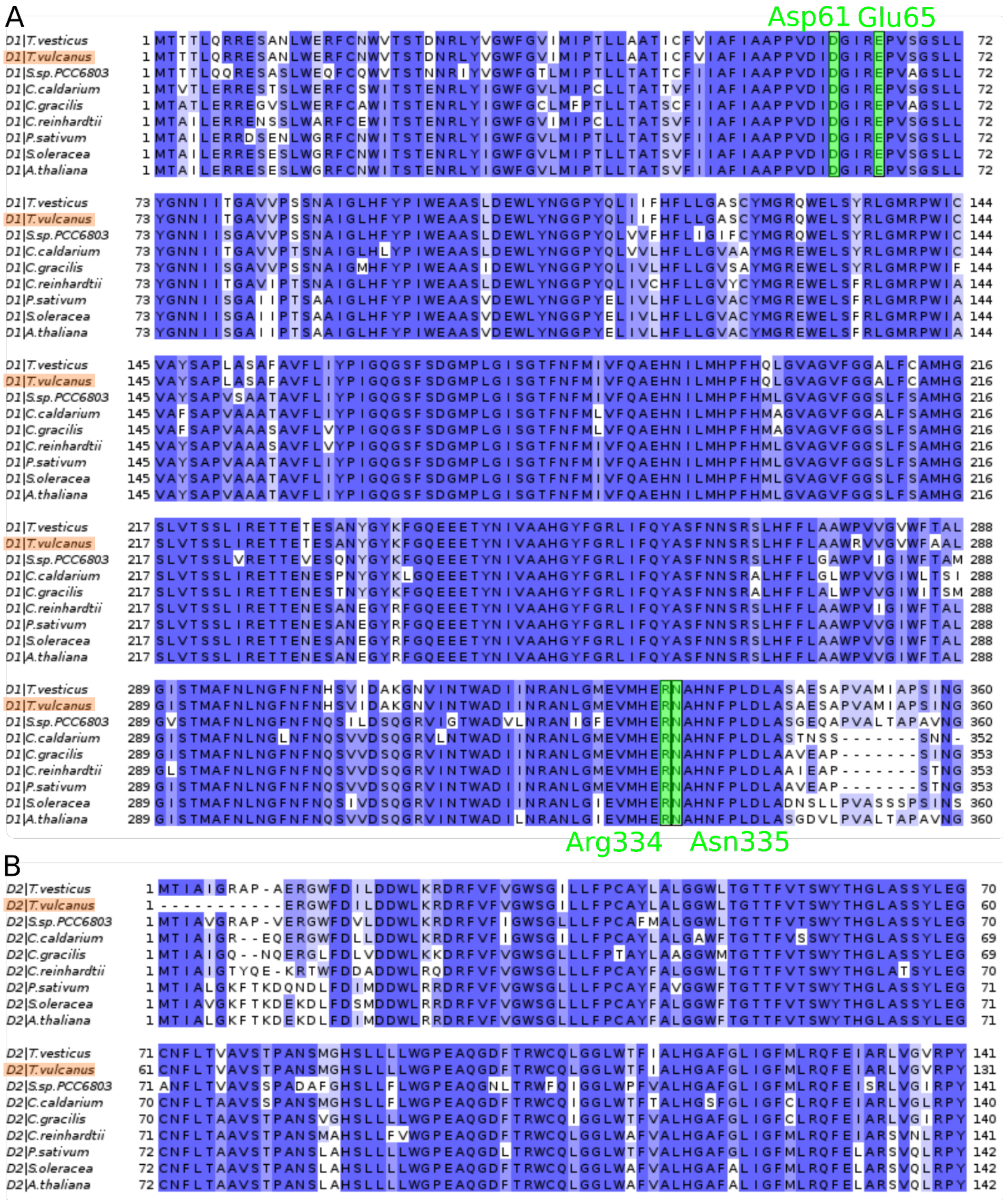


**Fig. S14. Multiple sequence alignment** of subunits **A)** D1, **B)** D2, and **C)** PsbO of PSII. Residues along the proposed proton transfer pathway in Cl1 are highlighted (similarities in green; differences in red). The residue numbering corresponds to *T. vulcanus* (in orange). All residues are highly conserved, with the exception of Arg115$_{PsbO}$ (Glu in *S.* sp. PCC 6803) and Arg184$_{PsbO}$ (Lys or Glu in other species). The sequences compared are: *T. vesticus* (7RF1)(2), *T. vulcanus* (3WU2)(1), *S.* sp. PCC 6803 (7N8O)(3), *C. caldarium* (4YUU)(4), *C. gracilis* (6JLU)(5), *C. reinhardtii* (6KAC)(6), *P. sativum* (5XNL)(7), *S. oleracea* (3JCU)(8), and *A. thaliana* (7OUI)(9). The sequence identity between *T. vulcanus* and the different species are 83-100% (D1), 85-100% (D2), and 41-82% (PsbO), respectively.

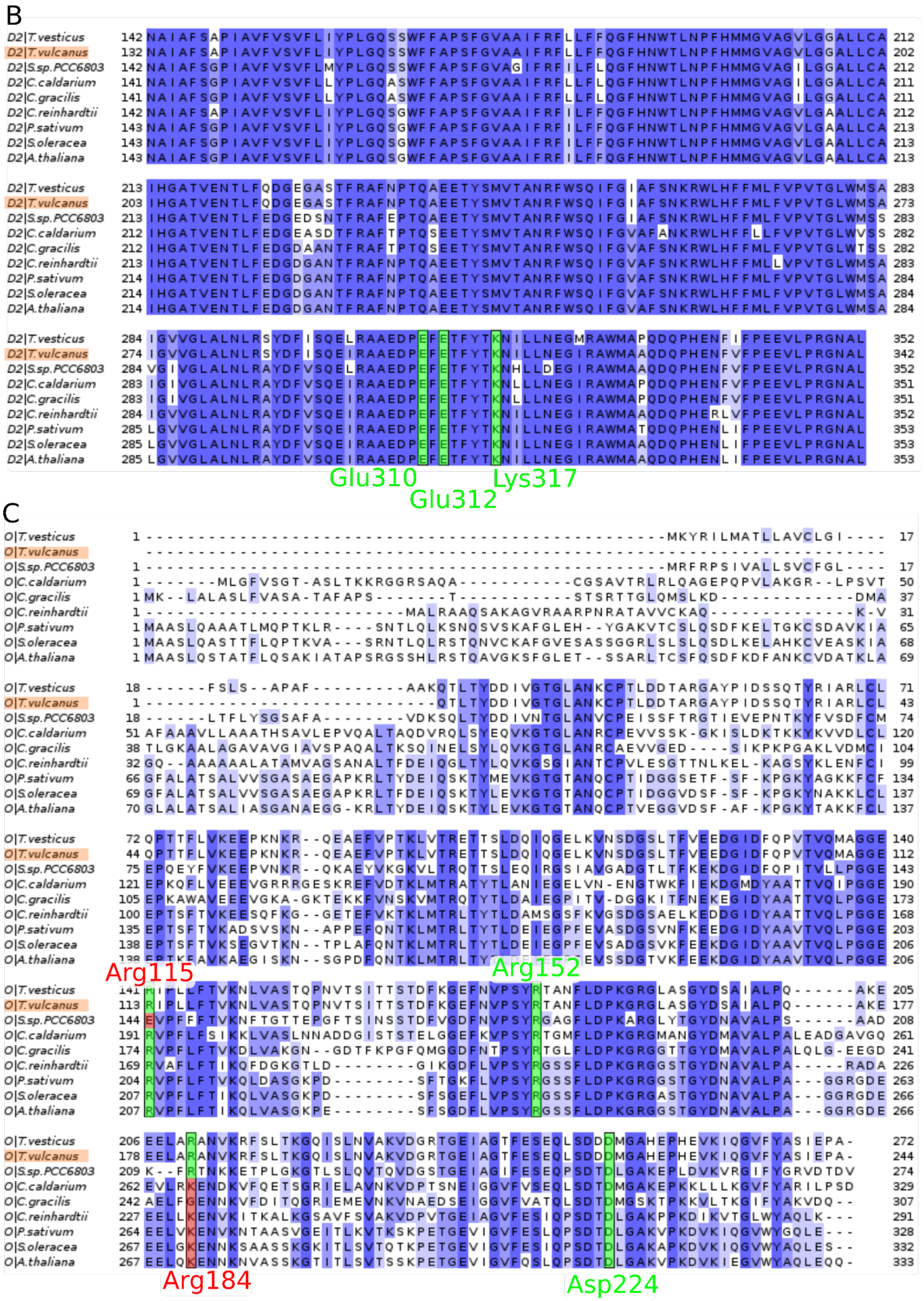


**Fig. S14 *(contd.)*. Multiple sequence alignment** of subunits **A**) D1, **B**) D2, and **C**) PsbO of PSII.

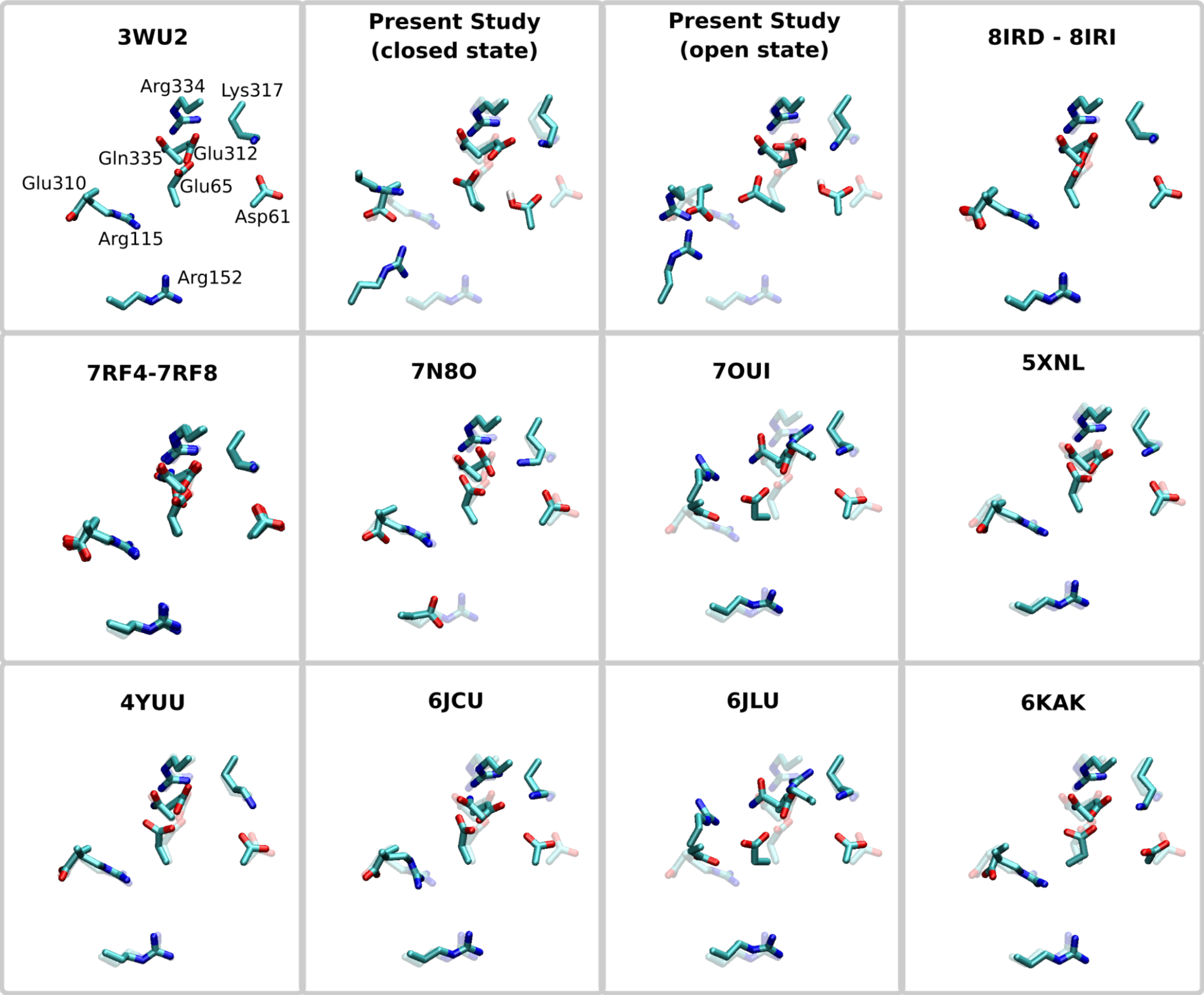


**Fig. S15. Structural comparison between predicted conformational changes and x-ray structures.** Representative structural snapshots of the open and the closed state of the Glu65 gate from present work, the dark state x-ray structure (PDB ID:3WU2 (1)) (transparent color), and XFEL structures (PDB ID:8IRD-8IRI (10), 7RF4-7RF8(2), 7N8O(3), 7OUI(9), 5XNL(7), 4YUU(4), 6JLU(5), 3JCU(8), and 6KAC(6)).

**Table S1. List of classical MD simulations,** with the modeled S states and protonated carboxylate residue indicated.

| S state | Protonated residue | Replicas and simulation length |
|---|---|---|
| $S_2Y_z^{\bullet}$ | Asp61 | 3 x 200 ns |
| | Glu312 | 3x 200 ns |
| | Glu65 | 3 x 200 ns |
| | Glu310 | 3 x 200 ns |
| | Asp224 | 2 x 200 ns |
| $S_3Y_z$ | Asp61 | 3 x 200 ns |
| | Glu312 | 3 x 200 ns |
| | Glu65 | 3 x 200 ns |
| | Glu310 | 3 x 200 ns |
| | Asp224 | 2 x 200 ns |
| $S_3Y_z^{\bullet}$ | Asp61 | 3 x 200 ns |
| | Glu312 | 3 x 200 ns |
| | Glu65 | 3 x 200 ns |
| | Glu310 | 3 x 200 ns |
| | Asp224 | 2 x 200ns |
| $S_4Y_z$ | Asp61 | 3 x 200 ns |
| | Glu312 | 3 x 200 ns |
| | Glu65 | 3 x 200 ns |
| | Glu310 | 3 x 200 ns |
| | Asp224 | 2 x 200 ns |
| | **Total:** | 10.4 µs |

**Table S2. List of QM/MM-MD simulations** and modeled protonated residues along the proton transfer chain.

| S state | Protonated residue(s) | Replicas and simulation length |
|---|---|---|
| $S_3Y_z$ | Asp61 | 2 x 60 ps |
| | Glu312 | 2 x 50 ps |
| | Glu65 | 2 x 50 ps |
| | Glu310 | 2 x 50 ps |
| | Asp224 | 1 x 50 ps |
| | Asp61 and Glu312 | 2 x 35 ps |
| | Asp61 and Glu65 | 2 x 35 ps |
| | Glu312 and Glu65 | 2 x 4 ps |
| $S_3Y_z$ * | Asp61 | 2 x 80 ps |
| | Glu312 | 2 x 50 ps |
| | Glu65 | 2 x 50 ps |
| | Glu310 | 2 x 50 ps |
| | Asp224 | 1 x 50 ps |
| | Asp61 and Glu312 | 2 x 35 ps |
| | Asp61 and Glu65 | 2 x 35 ps |
| | Glu312 and Glu65 | 2 x 4 ps |
| | Asp61* | 2 x 75 ps |
| | **Total:** | 1400 ps (1.4 ns) |

* based on $S_3Y_z$ MD snapshot

**Table S3. List of QM/MM free energy simulations** (QM/MM-US).

| S state | Proton transfer reaction | Protonated residue | Simulation length |
|---|---|---|---|
| $S_3Y_z$ | Asp61→ Glu312 | X | 15 x 4 ps |
| | | Glu65 | 15 x 2 ps |
| | Glu312 → Glu65 | X | 7 x 4 ps |
| | | Asp61 | 7 x 2 ps |
| | Glu65 → Glu310 | X | 11 x 4 ps |
| | | Glu312 | 12 x 2 ps |
| | Glu310 → Asp224 | X | 18 x 4 ps |
| $S_3Y_z^{\bullet}$ | Asp61→ Glu65 | X | 13 x 4 ps |
| | | Glu312 | 14 x 2 ps |
| | Asp61 → Glu312 | X | 13 x 4 ps |
| | | Glu65 | 16 x 2 ps |
| | Glu312 → Glu65 | X | 7 x 4 ps |
| | | Asp61 | 7 x 2 ps |
| | Glu65 → Glu310 | X | 13 x 4 ps |
| | | Glu312 | 13 x 2 ps |
| | Glu310 → Asp224 | X | 19 x 4 ps |
| | **Total:** | | 632 ps (0.6 ns) |
